# Sympathetic innervation of interscapular brown adipose tissue is not a predominant mediator of OT-elicited reductions of body weight gain and adiposity in male diet-induced obese rats

**DOI:** 10.1101/2024.09.12.612710

**Authors:** Melise M. Edwards, Ha K. Nguyen, Andrew D. Dodson, Adam J. Herbertson, Mackenzie K. Honeycutt, Jared D. Slattery, June R. Rambousek, Edison Tsui, Tami Wolden-Hanson, Tomasz A. Wietecha, James L. Graham, Geronimo P. Tapia, Carl L. Sikkema, Kevin D. O’Brien, Thomas O. Mundinger, Elaine R. Peskind, Vitaly Ryu, Peter J. Havel, Arshad M. Khan, Gerald J. Taborsky, James E. Blevins

## Abstract

Recent studies indicate that central administration of oxytocin (OT) reduces body weight (BW) in high fat diet-induced obese (DIO) rodents by reducing energy intake and increasing energy expenditure (EE). Previous studies in our lab have shown that administration of OT into the fourth ventricle (4V; hindbrain) elicits weight loss and stimulates interscapular brown adipose tissue temperature (T_IBAT_) in DIO rats. We hypothesized that OT-elicited stimulation of sympathetic nervous system (SNS) activation of IBAT contributes to its ability to activate BAT and reduce BW in DIO rats. To test this, we determined the effect of disrupting SNS activation of IBAT on OT-elicited stimulation of T_IBAT_ and reduction of BW in DIO rats. We first confirmed that bilateral surgical SNS denervation to IBAT was successful based on having achieved ≥ 60% reduction in IBAT norepinephrine (NE) content from DIO rats. NE content was selectively reduced in IBAT by 94.7 ± 2.7, 96.8 ± 1.8 and 85.9 ± 6.1% (*P*<0.05) at 1, 6 and 7-weeks post-denervation, respectively, and was unchanged in liver or inguinal white adipose tissue. We then measured the impact of bilateral surgical SNS denervation to IBAT on the ability of acute 4V OT (1, 5 µg) to stimulate T_IBAT_ in DIO rats. We found that the high dose of 4V OT (5 µg) stimulated T_IBAT_ similarly between sham and denervated rats (*P*=NS) and that the effects of 4V OT to stimulate T_IBAT_ did not require beta-3 adrenergic receptor signaling. We subsequently measured the effect of bilateral surgical denervation of IBAT on the effect of chronic 4V OT (16 nmol/day) or vehicle infusion to reduce BW, adiposity, and energy intake in DIO rats. Chronic 4V OT reduced BW gain by –7.2 ± 9.6 g and –14.1 ± 8.8 g in sham and denervated rats (*P*<0.05 vs vehicle treatment), respectively, and this effect was similar between groups (*P*=NS). These effects were associated with reductions in adiposity and energy intake (*P*<0.05). Collectively, these findings support the hypothesis that sympathetic innervation of IBAT is not required for central OT to increase BAT thermogenesis and reduce BW gain and adiposity in male DIO rats.

## Introduction

While the pleiotropic hormone, oxytocin (OT), is largely associated with male and female sexual and reproductive behavior (i.e. uterine contraction, milk ejection reflex, and penile erection) [1; 2] and social behaviors (i.e. parent-infant bonds, pair bonds, and increased trust) [3; 4], questions remain as far as how OT functions in the regulation of body weight (BW) [5; 6; 7; 8]. Although suppression of energy intake is thought to contribute to OT-elicited weight loss, pair-feeding studies in mice and rats suggest that OT’s effects on weight loss cannot be completely explained by its ability to reduce energy intake [9; 10; 11].

In addition to OT’s well-characterized effects on energy intake, previous studies have shown that OT increases other metabolic functions that control BW, namely energy expenditure (EE) [12; 13; 14; 15]. While brown adipose tissue thermogenesis (BAT) is important in the control of EE (see [16; 17] for review), it is unclear if OT’s effects on EE stem from 1) non-shivering BAT thermogenesis, 2) non-shivering and shivering thermogenesis in skeletal muscle [18], 3) spontaneous physical activity-induced thermogenesis [19], 4) white adipose tissue thermogenesis or 5) hormonal mediators (e.g. leptin [20], secretin [21; 22], irisin [23], fibroblast growth factor-21 [24] or thyroid hormone [25] (see [26; 27] for review). We have found that acute CNS injections of OT increase interscapular BAT temperature (T_IBAT_) (functional readout of BAT thermogenesis) in mice and rats [28; 29]. Moreover, the timing of OT-elicited elevations of T_IBAT_ coincide with the onset of OT-elicited reductions of BW in diet-induced obese (DIO) rats [28]. Sutton and colleagues reported that chemogenetic activation of OT neurons within the paraventricular nucleus (PVN) stimulates both subcutaneous BAT temperature and EE in *Oxytocin-Ires-Cre* mice [30]. In addition, Yuan et al found that systemic administration of OT increases thermogenic gene expression in interscapular brown adipose tissue (IBAT) and stimulates BAT differentiation *in vitro* in DIO mice [31]. In contrast, genetic or pharmacologically-induced deficits in OT signaling are associated with reductions in BAT thermogenesis [32; 33; 34; 35], EE [14; 15; 32; 36] and obesity [15; 36; 37; 38] in mice. More recently, we found that SNS innervation of IBAT is not a predominant mediator of OT-elicited reductions of BW and adiposity in male diet-induced obese mice [29]. One outstanding question is whether OT-elicited elevation of IBAT thermogenesis and weight loss require increased sympathetic nervous system (SNS) outflow to IBAT in DIO rats and whether the effect of OT on BAT thermogenesis may involve beta-3 adrenergic receptors (β3-AR) and hindbrain OTR. Here, we sought to clarify the role of SNS outflow to IBAT in mediating the effects of hindbrain OTR stimulation on both weight loss and BAT thermogenesis in a DIO rat model. Furthermore, we measured gross motor activity to test the extent to which increases in spontaneous physical activity may contribute to the effects of acute 4V OT on BAT thermogenesis.

We hypothesized that OT-elicited stimulation of sympathetic nervous system (SNS) activation of IBAT contributes to its ability to activate BAT and reduce BW in DIO rats. We initially verified the success of the IBAT denervation procedure by measuring IBAT norepinephrine (NE) content at 1-week post-denervation in lean rats. We then determined whether these effects can be replicated in DIO rats at 1, 6 and 7-weeks post-sham/denervation. To assess the role of SNS innervation of BAT in contributing to OT-elicited increases in BAT thermogenesis, we determined the impact of bilateral surgical SNS denervation to IBAT on the ability of acute 4V OT (1, 5 μg) to stimulate T_IBAT_ (as functional measure of BAT thermogenesis) in DIO rats. To assess the role of SNS innervation of BAT in contributing to OT-elicited weight loss, we subsequently measured the effect of bilateral surgical denervation of IBAT on the effect of chronic 4V OT (16 nmol/day over 29 days) or vehicle infusion to reduce BW, adiposity, and energy intake in DIO rats. We subsequently examined whether these effects were associated with a decrease adipocyte size and energy intake. In addition, we measured the temporal profile of acute 4V OT on BAT temperature, core temperature and gross motor activity to determine if spontaneous physical activity might have contributed to the effects of 4V OT on BAT thermogenesis. Lastly, we determined the extent to which 4V OT activates an early marker of neuronal activation, phosphorylated extracellular signal-regulated kinases 1 and 2-immunoreactivity (pERK1/2), within the hindbrain NTS in DIO rats.

## Methods

### Animals

Adult male Long-Evans rats (∼ 326–932 g BW) were initially obtained from Envigo (Indianapolis, IN) and maintained for at least 4 months on a high fat diet (HFD) prior to study onset. All animals were housed individually in Plexiglas cages in a temperature-controlled room (22±2°C) under a 12:12-h light-dark cycle. All rats were maintained on a 6 a.m./6 p.m. light cycle except for those in **Study 7A** [(7 a.m./7 p.m. light cycle (telemetry studies) or 1 a.m./1 p.m. reversed light cycle (energy intake studies)] and **Study 7B** (1 a.m./1 p.m. reversed light cycle). Rats had *ad libitum* access to water and a HFD providing 60% kcal from fat (Research Diets, D12492, New Brunswick, NJ). By design, rats used in **Study 1** and **Supplemental Study 1** were maintained on a low-fat chow diet (13% kcal from fat; diet no. 5001, LabDiet, St. Louis, MO). The research protocols were approved both by the Institutional Animal Care and Use Committee of the Veterans Affairs Puget Sound Health Care System (VAPSHCS) and the University of Washington in accordance with NIH’s *Guide for the Care and Use of Laboratory Animals* (NAS, 2011) [39] .

### Drug Preparation

Fresh solutions of OT acetate salt (Bachem Americas, Inc., Torrance, CA) were prepared the day of each experiment. OT was solubilized in sterile water (**Studies 5 and 6**). Fresh solutions of OT acetate salt (Bachem Americas, Inc., Torrance, CA) were solubilized in sterile water, loaded into Alzet® minipumps (model 2004; DURECT Corporation, Cupertino, CA) and subsequently primed in sterile 0.9% saline at 37° C for approximately 40 hours prior to minipump implantation based on manufacturer’s recommended instructions (**Study 6**). The β3-AR agonist, CL 316243 (Tocris/Bio-Techne Corporation, Minneapolis, MN), was solubilized in sterile water each day of each experiment (**Study 3; Supplemental Study 2**). The β3-AR antagonist, SR 59230A, was dissolved in 100% dimethyl sulfoxide (DMSO) each day of each experiment (**Study 5**).

### SNS denervation procedure

The methods used to surgically denervated IBAT in rodents have been described previously [29]. A dissecting microscope (Leica M60/M80; Leica Microsystems, Buffalo Grove, IL) was used throughout the procedure. A 1” midline incision was made in the skin dorsally at the level of the thorax and continued rostrally to the base of the skull. Connective tissue was blunt-dissected away from the adipose tissue with care to avoid cutting the large thoracodorsal artery that is located medially to both pads. Both left and right fat pads were separated from the midline. Each IBAT pad was lifted up and the intercostal nerve bundles were located below. Once the nerves were located, a sharp-point forceps was used to pull the nerve bundles straight up while using a 45 degree scissors to cut and remove 3-5 mm of nerves. The interscapular incision was closed with 4-0 non-absorbable monofilament Ethilon (nylon) sutures or with standard metal wound clips. Nerves were similarly identified but not cut for sham-operated animals. Rats were treated pre-operatively with the analgesic ketoprofen (2 mg/kg; Fort Dodge Animal Health, Overland Park, KS) prior to the completion of the denervation or sham procedure. This procedure was combined with transponder implantations for studies that involved IBAT temperature measurements in response to acute 4V injections (**Study 4**). Animals were allowed to recover for approximately 5–7 days prior to implantation of 4V cannulas.

### 4V cannulations for acute injections

The methods used to implant a cannula directed towards the 4V for acute 4V injections have been described previously [40; 41]. Animals were implanted with a cannula (P1 Technologies, Roanoke, VA) that was directed towards the 4V as previously described [42; 43; 44]. Briefly, rats under isoflurane anesthesia were placed in a stereotaxic apparatus [Digital Lab Standard Stereotaxic, Rat, (Item 51900), Stoelting Co., Wood Dale, IL] with the incisor bar positioned 3.3 mm below the interaural line. A 26-gauge cannula (P1 Technologies) was stereotaxically positioned into the 4V [–3.5 mm caudal to the interaural line; 1.4 mm lateral to the midline, and 6.2 mm ventral to the skull surface [45]] and secured to the surface of the skull with dental cement and stainless-steel screws.

### 4V cannulations for chronic infusions

The methods used to implant a cannula into the 4V for chronic infusions have been described previously [40; 41]. Briefly, rats were implanted with a cannula within the 4V with a side port that was connected to an osmotic minipump (model 2004, DURECT Corporation, Cupertino, CA) as previously described [28; 46]. Rats under isoflurane anesthesia were placed in a stereotaxic apparatus [Digital Lab Standard Stereotaxic, Rat, (Item 51900) Stoelting Co.] with the incisor bar positioned 3.3 mm below the interaural line. A 30-gauge cannula (P1 Technologies) was stereotaxically positioned into the 4V [–3.5 mm caudal to the interaural line; 1.4 mm lateral to the midline, and 7.2 mm ventral to the skull surface [45]] and secured to the surface of the skull with dental cement and stainless steel screws. Rats were treated with the analgesic ketoprofen (2 mg/kg; Fort Dodge Animal Health) and the antibiotic enrofloxacin (5 mg/kg; Bayer Healthcare LLC., Animal Health Division, Shawnee Mission, KS) at the completion of the 4V cannulations and were allowed to recover at least 10 days prior to implantation of osmotic minipumps.

### Implantation of implantable telemetry devices (PDT-4000 E-Mitter) into abdominal cavity

Animals were anesthetized with isoflurane and subsequently underwent a sham surgery (no implantation) or received implantations of a sterile PDT-4000 E-Mitter (23 mm long × 8 mm wide; Starr Life Sciences Company) into the intraperitoneal cavity. The abdominal opening was closed with 4-0 Vicryl absorbable suture and the skin was closed with 4-0 monofilament nonabsorbable suture. Vetbond glue was used to seal the wound and bind any tissue together between the sutures. Sutures were removed within two weeks after the PDT-4000 E-Mitter implantation. All PDT-4000 E-Mitters were confirmed to have remained within the abdominal cavity at the conclusion of the study.

### Implantation of temperature transponders underneath IBAT

The methods used for implanting a temperature transponder underneath IBAT in rats have been described previously [41]. Animals were anesthetized with isoflurane and had the dorsal surface along the upper midline of the back shaved and the area was scrubbed with 70% ethanol followed by betadine swabs. A one-inch incision was made at the midline of the interscapular area. The temperature transponder (14 mm long/2 mm wide) (HTEC IPTT-300; Bio Medic Data Systems, Inc., Seaford, DE) was implanted underneath the left IBAT pad as previously described [28; 47; 48]

### Acute IP or 4V injections and measurements of T_IBAT_

The methods used for IP or 4V injections and measurements of T_IBAT_ have been described previously [41]. On an experimental day, animals received either IP (CL 316243 or saline vehicle; 0.1 ml/kg injection volume) or 4V injections (OT or saline vehicle; 1 μL injection volume, 4V) during the early part of the light cycle following 4 hours of food deprivation. Injections were completed in a crossover design over approximately 7-day (CL 316243) or 48-h (OT) intervals such that each animal served as its own control. Animals remained without access to food for an additional 4 h (**Studies 3 & 4**) during the T_IBAT_ measurements. A handheld reader (DAS-8007-IUS Reader System; Bio Medic Data Systems, Inc.) was used to collect measurements of T_IBAT_.

### Changes of Core Temperature and Gross Motor Activity

Telemetry recordings of core temperature and gross motor activity were measured from each rat in the home cage immediately prior to injections and for a 4-h period after injections. Core temperature and gross motor activity were recorded every 5 min.

### Body Composition

Determinations of lean body mass and fat mass were made on un-anesthetized rats by quantitative magnetic resonance using an EchoMRI 4-in-1-700^TM^ instrument (Echo Medical Systems, Houston, TX) at the VAPSHCS Rodent Metabolic Phenotyping Core. Measurements were taken prior to 4V cannulations and minipump implantations as well as at the end of the infusion period.

### Tissue collection for Norepinephrine (NE) content measurements

Rats were euthanized by rapid conscious decapitation at 1 week (**Study 1**), 6 weeks (**Study 2**), 7 weeks (**Study 2, Study 5**), 9 weeks (**Study 6**) or 10–11 weeks (**Study 4**) post-sham or denervation procedure. Trunk blood and tissues (IBAT, EWAT, IWAT, liver and/or pancreas) were collected from 4-h fasted rats. Tissue was rapidly removed, wrapped in foil and frozen in liquid N_2_. Samples were stored frozen at –80°C until analysis. Note that anesthesia was not used when collecting tissue for NE content as it can cause the release of NE from SNS terminals within the tissue [49].

### NE content measurements (biochemical confirmation of IBAT denervation procedure)

NE content was measured in IBAT, EWAT, IWAT, liver and/or pancreas using previously established techniques [50]. Successful denervation was noted by ≥ 60% reduction in IBAT NE content as previously noted [51]. Experimental animals that did not meet this criterion were excluded from the data analysis.

## Study Protocols

### Study 1A: Determine the success of the surgical denervation of only the superficial nerves at 1-week post-sham or denervation in lean rats by measuring NE content

Rats underwent sham or bilateral SNS denervation procedures of only the superficial nerves that innervate IBAT and, to prevent the confound of anesthesia on NE content, animals were euthanized by rapid conscious decapitation at 1-week post-sham or denervation procedure.

### Study 1B: Determine the success of the surgical denervation procedure at 1-week post-sham or denervation in lean rats by measuring NE content

Rats underwent sham or bilateral SNS denervation procedures of both superficial and more ventrally located intercostal nerves that innervate IBAT and, to prevent the confound of anesthesia on NE content, animals were euthanized by rapid conscious decapitation at 1-week post-sham or denervation procedure.

### Study 2: Determine the success of the surgical denervation procedure at 1, 6 and 7-weeks post-sham or denervation in DIO rats by measuring NE content

Rats were fed *ad libitum* and maintained on HFD for approximately 5 months prior to underdoing sham or SNS denervation procedures. In addition to weekly BW measurements, body composition measurements were obtained at baseline and at 6 and 7-weeks post-denervation/sham procedures. Rats were euthanized by rapid conscious decapitation 1, 6 and 7-weeks post-sham or denervation procedure.

### Study 3: Determine if surgical denervation of IBAT changes the ability of the β-3R agonist, CL 316243, to increase T_IBAT_ in DIO rats

Rats were fed *ad libitum* and maintained on HFD for approximately 4.25 months prior to underdoing sham or SNS denervation procedures and implantation of temperature transponders underneath the left IBAT depot. Rats were allowed to recover for at least 1 week during which time they were adapted to a daily 4-h fast, handling, and mock injections. On an experimental day, 4-h fasted rats received CL 316243 (0.1 or 1 mg/kg) or vehicle (sterile water) during the early part of the light cycle in a crossover design at approximately 7-day intervals such that each animal served as its own control (approximately 1–3 weeks post-sham or denervation procedures). T_IBAT_ was measured at baseline (–2 h; 9:00 a.m.), immediately prior to IP injections (0 h; 9:45–10:00 a.m.), and at 0.25, 0.5, 0.75, 1, 1.25, 1.5, 2, 3, 4, and 24-h post-injection (10:00 a.m.). Energy intake and BW were measured daily. This dose range was based on doses of CL 316243 found to be effective at reducing energy intake and weight gain in rats [52; 53]. Animals were euthanized by rapid conscious decapitation at 13 weeks post-sham or denervation procedure.

### Study 4: Determine the extent to which OT-induced activation of sympathetic outflow to IBAT contributes to its ability to increase T_IBAT_ in DIO rats

Rats from **Study 3** were implanted with 4V cannulas approximately 1 month following sham/denervation procedures and implantation of temperature transponders underneath the left IBAT depot. Rats were allowed to recover for at least 2 weeks during which time they were adapted to a daily 4-h fast, handling, and mock injections. On an experimental day, 4-h fasted rats received OT (1 or 5 μg/μl) or vehicle during the early part of the light cycle in order to maximize the effects of OT [15; 53] during a time when circulating NE levels [54] and IBAT catecholamine levels are lower [55]. Injections were completed in a crossover design at approximately 48-h intervals such that each animal served as its own control (approximately 7-weeks post-sham or denervation procedures). T_IBAT_ was measured at baseline (–2 h; 9:00 a.m.), immediately prior to 4V injections (0 h; 9:45– 10:00 a.m.), and at 0.25, 0.5, 0.75, 1, 1.25, 1.5, 2, 3, 4, and 24-h post-injection (10:00 a.m.). Energy intake and BW were measured daily. This dose range was based on doses of 4V OT found to be effective at stimulating T_IBAT_ in DIO rats in previous studies [28].

### Study 5: Determine the extent to which OT-induced activation of sympathetic outflow to IBAT requires activation of β3-AR to increase T_IBAT_ in DIO rats

Rats were fed *ad libitum* and maintained on HFD for approximately 4 months prior to being implanted with temperature transponders underneath the left IBAT depot. Rats were subsequently implanted with 4V cannulas approximately 1 month following implantation of temperature transponders. Rats were allowed to recover for 4 weeks during which time they were adapted to a daily 4-h fast, handling, and mock injections. On an experimental day, 4-h fasted rats received the β3-AR antagonist, SR 59230A (0.5 mg/kg, IP) or vehicle (DMSO) approximately 20 min prior to 4V administration of either OT (5 μg/μl) or vehicle. Injections occurred during the early part of the light cycle in order to maximize the effects of OT [15; 53] during a time when circulating NE levels [54] and IBAT catecholamine levels are lower [55]. Injections were completed in a crossover design at approximately 7-day intervals such that each animal served as its own control. T_IBAT_ was measured at baseline (-2 h; 9:00 a.m.), immediately prior to 4V injections (0 h; 9:45-10:00 a.m.), and at 0.25, 0.5, 0.75, 1, 1.25, 1.5, 2, 3, 4, and 24-h post-injection (10:00 a.m.). Energy intake and BW were measured daily. This dose of SR 59230A was based on other studies [56; 57; 58] and doses that we found to be subthreshold for reducing T_IBAT_ in our pilot studies (data not shown) and able to block the effects of the β3-AR agonist, CL 316243, on T_IBAT_ in DIO rats (**Supplemental Figure 1**).

### Study 6A: Determine the extent to which OT-induced activation of sympathetic outflow to IBAT contributes to its ability to elicit weight loss in DIO rats

Rats were fed *ad libitum* and maintained on HFD for at least 4.5 months prior to receiving 4V cannulas and minipumps to infuse vehicle or OT (16 nmol/day) over 29 days as previously described [28]. This dose was selected based on a dose of 4V OT found to be effective at reducing BW in DIO rats [28]. Daily energy intake and BW were also tracked for 29 days. Animals were euthanized by rapid conscious decapitation at 7 weeks post-sham or denervation procedure. Trunk blood and tissues [IBAT, epididymal white adipose tissue (EWAT), inguinal white adipose tissue (IWAT), liver and pancreas)] were collected from 4-h fasted rats and tissues were subsequently analyzed for IBAT NE content to confirm success of denervation procedure relative to sham-operated animals and other tissues (EWAT, IWAT, liver and pancreas).

### Study 6B: Determine the success of the surgical denervation procedure at 6-8 weeks post-sham or denervation in DIO rats by measuring thermogenic gene expression

Rats from **Study 6A** were used for these studies. All rats underwent sham or bilateral surgical SNS denervation procedures prior to being implanted with 4V cannulas and minipumps. Animals were euthanized by rapid conscious decapitation at approximately 6–8 weeks post-sham or denervation procedure. Tissues were collected following a 4-h fast.

### Study 6C: Determine the extent to which OT-induced activation of sympathetic outflow to IBAT impacts thermogenic gene expression in IBAT and EWAT in DIO mice

Rats from **Study 6A** were used for these studies. All rats underwent sham or bilateral surgical SNS denervation procedures prior to being implanted with 4V cannulas and minipumps. Animals were euthanized by rapid conscious decapitation at approximately 6–8 weeks post-sham or denervation procedure. Tissues were collected following a 4-h fast.

### Study 7A: Determine the extent to which 4V OT increases gross motor activity, core temperature, T_IBAT_ and energy intake in DIO rats

#### Telemetry measurements

Rats were fed *ad libitum* and maintained on HFD for approximately 6 months prior to being implanted with temperature transponders underneath IBAT, intra-abdominal telemetry devices and 4V cannulas as previously described. Rats were implanted temperature transponders approximately 4 days prior to being implanted with intra-abdominal telemetry devices. Animals were allowed up to 3 weeks post-op recovery prior to being implanted with 4V cannulas. Rats were allowed to recover for at least 2 weeks during which time they were adapted to a daily 4-h fast, handling, and mock injections. Like **Study 4**, on an experimental day, 4-h fasted rats received OT (1 or 5 μg/μl) or vehicle during the early part of the light cycle. Injections were completed in a crossover design at approximately 48-h intervals such that each animal served as its own control. T_IBAT_ was measured at baseline (-2 h; 9:00 a.m.), immediately prior to 4V injections (0 h; 9:45-10:00 a.m.), and at 0.25, 0.5, 0.75, 1, 1.25, 1.5, 2, 3, 4, and 24-h post-injection (10:00 a.m.). Energy intake and BW were measured daily.

#### Energy intake measurements

Animals were treated identically as described above with the exception that animals were adapted to a 1 a.m. (lights on)/1 p.m. (lights off) light cycle and animals received acute 4V injections at the end of the light cycle (12:40-1 p.m.). Food was returned at the onset of the dark cycle. Food was measured at 0.5, 1, 2, 3, 4 and 24-h post-injection.

### Study 7B: Determine the extent to which 4V OT activates an early marker of neuronal activation, phosphorylated extracellular signal-regulated kinases 1 and 2-immunoreactivity (pERK1/2), within the hindbrain NTS in DIO rats

#### Atlas nomenclature

We used the standardized nomenclature system provided by Swanson (2018) [56] to describe and name regions of the hindbrain for which we mapped activation patterns. This approach is being used by us as part of our longstanding effort to encourage investigators to place their data into a common spatial model so that they are interrelated with other datasets mapped in the same space [59; 60]. This also requires a common nomenclature. First devised in a comprehensive reference work for human brain [61], the nomenclature system includes an *(author, date)* attribution as *part of the name of the structure*, to help readers know the provenance of where the term was first described as defined in the atlas. For those terms where such information was not traced, the assignation “(>1840)” is used instead to note that it was in use sometime after the development of cell theory. We applied this nomenclature to those regions of the hindbrain we examined for the early activation marker, phosphorylated p44/42 MAP Kinases (pERK1/2). These regions include the *area postrema (>1840)* (AP), *nucleus of solitary tract (>1840)* (NTS), and *dorsal motor nucleus of vagus nerve (>1840)* (DMX).

#### Subjects

Rats from **Study 7A** were used for this study. Following the conclusion of **Study 7A**, 4-h fasted rats received OT (5 μg/μl) or vehicle during the late part of the light cycle. Animals remained fasted for an additional 0.25 or 0.5-h post-injection, at which time animals were euthanized with an IP overdose of ketamine cocktail and transcardially perfused with PBS followed by 4% paraformaldehyde in 0.1 M PBS. Brains were removed, stored overnight in fresh fixative at 4°C and subsequently transferred to 0.1 M PBS containing 25% sucrose for 48 h. Brains were then frozen by submersion for 20–30 s in isopentane chilled with dry ice.

#### Tissue Processing and Immunohistochemistry

Perfusion-fixed frozen brain specimens were shipped overnight in dry ice to The University of Texas at El Paso for histological analysis, but arrived thawed due to an unforeseen shipping delay beyond our control. Therefore, they were prepared to be re-frozen by placing them into a 20% sucrose cryoprotectant solution and stored at 4°C before processing. Phospho-ERK1/2 labeling in the dorsal vagal complex did not appear to be adversely affected in our peroxidase-based reactions and was found in the same cellular compartments previously noted by others [62; 63; 64].

Brains were coronally-cut into two tissue blocks, and the hindbrain portion was re-frozen with its rostral face down onto an ice-cold solid brass freezing stage (3.25 × 7.25 × 1.5 in; Brain Research Laboratories, Newton, MA, USA; cat #3488-RJ) modified to fit a modern ‘Naples’ object-holder and carrier of a sliding microtome (Reichert; Vienna, Austria; Nr. 15 156). Dry ice pellets in the stage troughs were used to keep the stage at freezing temperatures. The frozen blocks were cut into 30 µm-thick coronal sections, collected as six series into 24-well tissue culture plates filled with cryoprotectant (50% 0.05 M sodium phosphate buffer, 30% ethylene glycol, 20% glycerol; pH 7.4 at room temperature), and stored at −20°C before undergoing further processing.

Using Tris-buffered saline (TBS) (0.05 M; pH 7.4 at room temperature), sections were washed of cryoprotectant (5 washes for 5 minutes each; 5 ✕ 5) and were incubated in 0.007% phenylhydrazine solution for 20 minutes at room temperature. This was followed by 5 ✕ 5 TBS washes, then an incubation in blocking solution consisting of TBS containing 2% (v/v) normal donkey serum (catalog #S30-100ML; EMD-Millipore, Burlington, MA) and 0.1% (v/v) Triton X-100 for ∼1 hour at room temperature. The tissue was then reacted with an antibody solution consisting of blocking solution containing a primary antibody targeting the phosphorylated forms of ERKs 1 and 2 (pERK1/2; EC 2.7.11.24) (1:100 dilution; catalog #9101S; Cell Signaling Technology, Danvers, MA; RRID:AB_331646) for ∼12 hours at 4°C. This was followed by a 5 ✕ 5 wash period, then reaction of the sections with an affinity-purified donkey anti-rabbit IgG secondary antibody (1:500 dilution; catalog #711-065-152; Jackson ImmunoResearch, West Grove, PA; RRID:AB_2340593) for ∼5 hours at room temperature. The sections underwent another 5 ✕ 5 wash before incubation in an avidin-biotin complex (ABC) reagent (VECTASTAIN® ABC-HRP Kit, catalog #PK-4000; Vector Laboratories, Newark, CA; RRID:AB_2336818) for ∼1 hour at room temperature. The tissue was rinsed via a 5 ✕ 5 TBS wash before being subjected to a Ni^2+^-enhanced 3,3’-diaminobenzidene peroxidase reaction to visualize antibody labeling.

Following the reaction, the sections underwent a final 5 ✕ 5 wash before being mounted onto gelatin subbed slides. Once air-dried, the mounted sections were dehydrated in ascending concentrations of ethanol (50%, 70%, 95%, 3 ✕ 100%; 3 min each), and cleared in xylenes (2 ✕ 12 min). As individual slides were removed from the staining jar, they were immediately coated with 5 drops of dibutyl phthalate xylenes (DPX) mountant for histology (cat #06522-100ML, EMD-Millipore, Burlington, MA), coverslipped, and laid to dry.

#### Wide-field Microscopy

Using an Olympus BX63 light microscope with an X–Y–Z motorized stage and an attached DP74 color CMOS camera (cooled, 20.8 MP pixel-shift, 60 fps; Olympus America, Inc., Waltham, MA), photomicrographs were captured under bright-field and dark-field illumination. Image acquisition was processed using cellSense™ Dimension software installed on a Hewlett Packard PC workstation. Wide-field mosaic photomicrographs of the sections were captured using a ×20 magnification objective lens (Olympus UPlanSApo, N.A. 0.50, FN 26.5 mm) and 15% overlap of stitched tiles. Each image was exported as full-resolution uncompressed TIFF-formatted files.

#### Anatomical observations for determining the plane of section

To assess the recruitment of ERK1/2 in the NTS of OT- and vehicle-treated animals, we relied on the *Brain Maps 4.0 (BM4.0)* open-access atlas [65] (RRID:SCR_017314) to localize sections to a particular region of the dorsal vagal complex (*BM4.0* atlas level 70) to have a stable landmark for comparing immunoreactive signal. This was performed by referencing the region delineations represented in *BM4.0* templates of caudal hindbrain levels, relying mostly on the morphology of the *area postrema (>1840)* (AP). The AP is represented at two *BM4.0* atlas levels, levels 69 (Bregma = −13.76 mm) and 70 (Bregma = −14.16 mm), respectively. At level 69, the dorsal AP boundaries extend laterally, and the nucleus has a more flattened dorsal border when compared to more caudal representations. Ventral to the *nucleus of solitary tract (>1840)* (NTS), the *dorsal motor nucleus of vagus nerve (>1840)* (DMX) is a large, blimp-shaped cell group that decreases in cellular area as the nucleus extends caudally. At level 70, the AP takes on a rounder shape and reduces in size as it, too, extends caudally along the dorsal midline. The cytoarchitectural features of these regions surrounding the NTS are reliable landmarks for the determination of a rostral or caudal AP-containing section and were used to select the sections that best corresponded to atlas level 70 for analysis.

#### Regional Parcellations and Cell Tabulation

Once the sections that most closely adhered to *BM4.0* atlas level 70 region boundaries were identified, the photomicrographs were imported into Adobe Illustrator CC (AI) (Version 28.6) and annotated manually. This was performed by superimposing the bright-field and dark-field section images over the *BM4.0* digital template. In a separate layer, the reviewer used the *Pencil Tool* to parcellate the region boundaries of the immunolabeled section, relying on the cytoarchitectural features to guide their tracings. The myeloarchitecture, that is more easily visualized under dark-field viewing conditions, was used to trace important white matter areas, such as the *solitary tract (>1840)* (ts). If a region was displaced due to damage of the tissue, then its inferred borders were represented.

Another layer was created to mark the locations of pERK1/2-immunoreactive cell profiles using the *Ellipse Tool*, in which the reviewer placed a colored ellipse over each immunolabeled perikaryon. Considering that the borders of the ventral NTS and dorsal DMX do not have a clear-cut boundary but rather contain an intermixing of cells, the morphology of each annotated cell profile was assessed for its consistency with the smaller neuronal size of the NTS or larger perikaryon of the motor DMX, so as to only consider activation of the NTS here.

#### Blood collection

Trunk blood was collected from 4-h (**Studies 1 & 2**, **Study 6A–C**), 4.25 or 4.5 h (**Study 7B**) or 6-h (**Studies 3, 4 & 5**) fasted rats within a 2-h window towards the beginning of the light cycle (10:00 a.m.–12:00 p.m.) as previously described in DIO CD^®^ IGS and Long-Evans rats and C57BL/6J mice [28; 43] with the exception of **Study 7B**, in which blood was collected towards the end of the light cycle. Treatment groups were counterbalanced at time of euthanasia to avoid time of day bias. Blood samples [up to 3 mL] were collected immediately prior to transcardial perfusion by cardiac puncture in chilled K2 EDTA Microtainer Tubes (Becton-Dickinson, Franklin Lakes, NJ). Whole blood was centrifuged at 6,000 rpm for 1.5 min at 4°C; plasma was removed, aliquoted and stored at –80°C for subsequent analysis.

#### Plasma hormone measurements

Plasma leptin and insulin were measured using electrochemiluminescence detection [Meso Scale Discovery (MSD^®^), Rockville, MD] using established procedures [28; 66]. Intra-assay coefficient of variation (CV) for leptin was 5.9% and 1.6% for insulin. The range of detectability for the leptin assay is 0.137–100 ng/mL and 0.069–50 ng/mL for insulin. Plasma fibroblast growth factor-21 (FGF-21) (R&D Systems, Minneapolis, MN) and irisin (AdipoGen, San Diego, CA) levels were determined by ELISA. The intra-assay CV for FGF-21 and irisin were 2.2% and 9.1%, respectively; the ranges of detectability were 31.3–2,000 pg/mL (FGF-21) and 0.078–5 μg/mL (irisin). Plasma adiponectin was also measured using electrochemiluminescence detection [Millipore Sigma (Burlington, MA)] using established procedures [28; 66]. The intra-assay CV for adiponectin was 3.7%. The range of detectability for the adiponectin assay is 2.8–178 ng/mL. The data were normalized to historical values using a pooled plasma quality control sample that was assayed in each plate.

#### Blood glucose and lipid measurements

Blood was collected for glucose measurements by tail vein nick in 4-h or 6-h fasted rats and measured with a glucometer using the AlphaTRAK 2 blood glucose monitoring system (Abbott Laboratories, Abbott Park, IL) [28; 67]. Total cholesterol (TC) [Fisher Diagnostics (Middletown, VA)] and free fatty acids (FFAs) [Wako Chemicals USA, Inc., Richmond, VA)] were measured using an enzymatic-based kits. Intra-assay CVs for TC and FFAs were 1.4 and 2.3%, respectively. These assay procedures have been validated for rodents [68].

#### Adipose tissue processing for adipocyte size

Adipose tissue depots were collected at the end of the infusion period in DIO rats from **Studies 6A–C** (IBAT, EWAT and IWAT). IBAT, IWAT, and EWAT depots were dissected and placed in 4% paraformaldehyde-PBS for 24 h and then placed in 70% ethanol (EtOH) prior to paraffin embedding. Sections (5 μm-thick) were obtained using a rotary microtome, slide-mounted using a floatation water bath (37°C) and baked for 30 min at 60°C to give approximately 15 or 16 slides/fat depot with two sections/slide.

#### Adipocyte size analysis

Adipocyte size analysis was performed on deparaffinized and digitized IWAT and EWAT sections. The average cell area from two randomized photomicrographs was determined using the built-in particle counting method of ImageJ software (National Institutes of Health, Bethesda, MD). Slides were visualized using bright field on an Olympus BX51 microscope (Olympus Corporation of the Americas; Center Valley, PA) and photographed using a Canon EOS 5D SR DSLR (Canon U.S.A., Inc., Melville, NY) camera at ×100 magnification. Values for each tissue within a treatment were averaged to obtain the mean of the treatment group.

#### Tissue collection for quantitative real-time PCR (qPCR)

Tissues from **Study 1A** (IBAT, IWAT, and liver), **Studies 1B–2** (IBAT, IWAT, liver and pancreas), **Studies 4 and 6** (IBAT, IWAT, EWAT, liver and pancreas) and **Study 7B** (brain) were collected from a subset of 4-h (**Studies 1A/B–2**, **Study 6**), 4.25 to 4.5-h fasted rats (**Study 7**), or 6-h fasted rats (**Study 4**). Tissues from **Studies 1-6** were collected within a 2-h window towards the start of the light cycle (10:00 a.m.–12:00 p.m.) as previously described in DIO CD^®^ IGS/Long-Evans rats and C57BL/6J mice [28; 43; 53]. Tissue (brain) from **Study 7B** was collected within a 2-h window towards the end of the light cycle. Tissue was rapidly removed, wrapped in foil and frozen in liquid N_2_. Samples were stored frozen at –80°C until analysis.

#### qPCR

RNA extracted from samples of IBAT and IWAT (**Studies 4–7**) were analyzed using the RNeasy Lipid Mini Kit (Qiagen Sciences Inc, Germantown, MD) followed by reverse transcription into cDNA using a high-capacity cDNA archive kit (Applied Biosystems, Foster City, CA). Quantitative analysis for relative levels of mRNA in the RNA extracts was measured in duplicate by qPCR on an Applied Biosystems 7500 Real-Time PCR system (Thermo Fisher Scientific, Waltham, MA) and normalized to the cycle threshold value of Nono mRNA in each sample. The TaqMan® probes used in the study were Thermo Fisher Scientific Gene Expression Assay probes. The probe for rat *Nono* (Rn01418995_g1), uncoupling protein-1 (UCP-1) (*Ucp1*; catalog no. Rn00562126_m1), uncoupling protein-2 (UCP-2) (*Ucp2*; catalog no. Mm00627599_m1), uncoupling protein-3 (UCP-3) (*Ucp3*; catalog no. Rn00565874_m1), β1-adrenergic receptor (β1-AR) (*Adrb1*; catalog no. Rn00824536_s1), β2-adrenergic receptor (β2-AR) (*Adrb2*; catalog no. Rn005600650_s1), β3-adrenergic receptor (β3-AR) (*Adrb3*; catalog no. Rn01478698_g1), alpha 2-adrenergic receptor (alpha 2-AR) (*Adra2a*; catalog no. Rn00562488_s1)], type 2 deiodinase (D2) (*Dio2*; catalog no. Rn00581867_m1), PR domain containing 16 (*Prdm16*; catalog no. Rn01516224_m1), cytochrome c oxidase subunit 8b (*Cox8b*; catalog no. Rn00562884_m1), G-protein coupled receptor 120 (*Gpr120*; catalog no. Rn01759772_m1), bone morphogenetic protein 8b (*bmp8b*; catalog no. Rn01516089_gH), cell death-inducing DNA fragmentation factor alpha-like effector A (*Cidea*; catalog no. Rn04181355_m1), peroxisome proliferator-activated receptor gamma coactivator 1 alpha (*Ppargc1a*; catalog no. Rn00580241_m1) and cannabinoid receptor 1(*Cnr1*; catalog no. Mm01212171_s1) were acquired from Thermo Fisher Scientific. Relative amounts of target mRNA were determined using the Comparative C_T_ or 2-^ΔΔC^_T_ method [69] following adjustment for the housekeeping gene, Nono. Specific mRNA levels of all genes of interest were normalized to the cycle threshold value of *Nono* mRNA in each sample and expressed as changes normalized to controls (vehicle/sham treatment).

#### Statistical Analyses

All results are expressed as means ± SE. Planned comparisons between multiple groups involving between-subjects designs were made using one-way ANOVA followed by a post-hoc Fisher’s least significant difference test. Two-way ANOVA was used to examine drug (OT vs vehicle)*surgery (sham vs denervation) interactive effects on body weight change and fat mass. Planned comparisons involving within-subjects designs were made using a one-way repeated-measures ANOVA followed by a post-hoc Fisher’s least significant difference test. Repeated measures ANOVA involving within-subjects designs were used to examine the effects of acute 4V OT on T_IBAT_, core temperature, activity, and energy intake across multiple time points. Two-way repeated measures ANOVA was used to examine the interactive effects of the either the β3-AR agonist, CL 316243, or 4V OT and the β3-AR antagonist, SR 59230A on T_IBAT_. Analyses were performed using the statistical program SYSTAT (Systat Software, Point Richmond, CA). Differences were considered significant at *P*<0.05, 2-tailed.

## Results

### Study 1A: Determine the success of the surgical denervation of only the superficial nerves at 1-week post-sham or denervation in lean rats by measuring NE content

Rats (N= 10 at study onset) were used for this study. The goal of this study was to determine the success of lesioning only the more superficial intercostal nerves that innervate IBAT in lean rats by confirming a reduction of NE content that was specific to IBAT relative to other tissues (IWAT and liver). All IBAT tissues from Study 1A animals were analyzed for IBAT NE content and 1 of the 5 animals was removed on account of having a failed IBAT denervation procedure. Lesions of only the more superficial nerves resulted in a tendency for IBAT NE content to be reduced in denervated rats relative to sham-operated control rats (80.3 ± 6.0%; *P*=0.118) (data not shown). NE content remained unchanged in IWAT or liver from denervated rats relative to sham rats (*P*=NS). There were also no significant differences in BWs between sham and denervation groups at 1-week post-sham/denervation surgery (*P*=NS; data not shown).

### Study 1B: Determine the success of the surgical denervation procedure at 1-week post-sham or denervation in lean rats by measuring NE content

Rats (N= 10 at study onset) were used for this study. The goal of this study was to verify the success of the SNS denervation of both superficial and more ventrally located intercostal nerves that innervate IBAT by confirming a reduction of NE content that was specific to IBAT relative to other tissues (IWAT, liver and pancreas). All IBAT tissues from **Study 1B** animals were analyzed for IBAT NE content and none of the 5 animals with IBAT denervation were removed on account of having a failed IBAT denervation procedure. IBAT NE content was reduced in denervated rats by 91.5±5.4% in denervated rats relative to sham-operated control rats [(F(1,8) = 57.941, *P*=0.000) (**Figure 1**). In contrast, NE content was unchanged in IWAT, liver or pancreas in denervated rats relative to sham rats (*P*=NS). There were no significant differences in BWs between sham and denervation groups at 1-week post-sham/denervation surgery (*P*=NS; data not shown).

**Figure 1.**
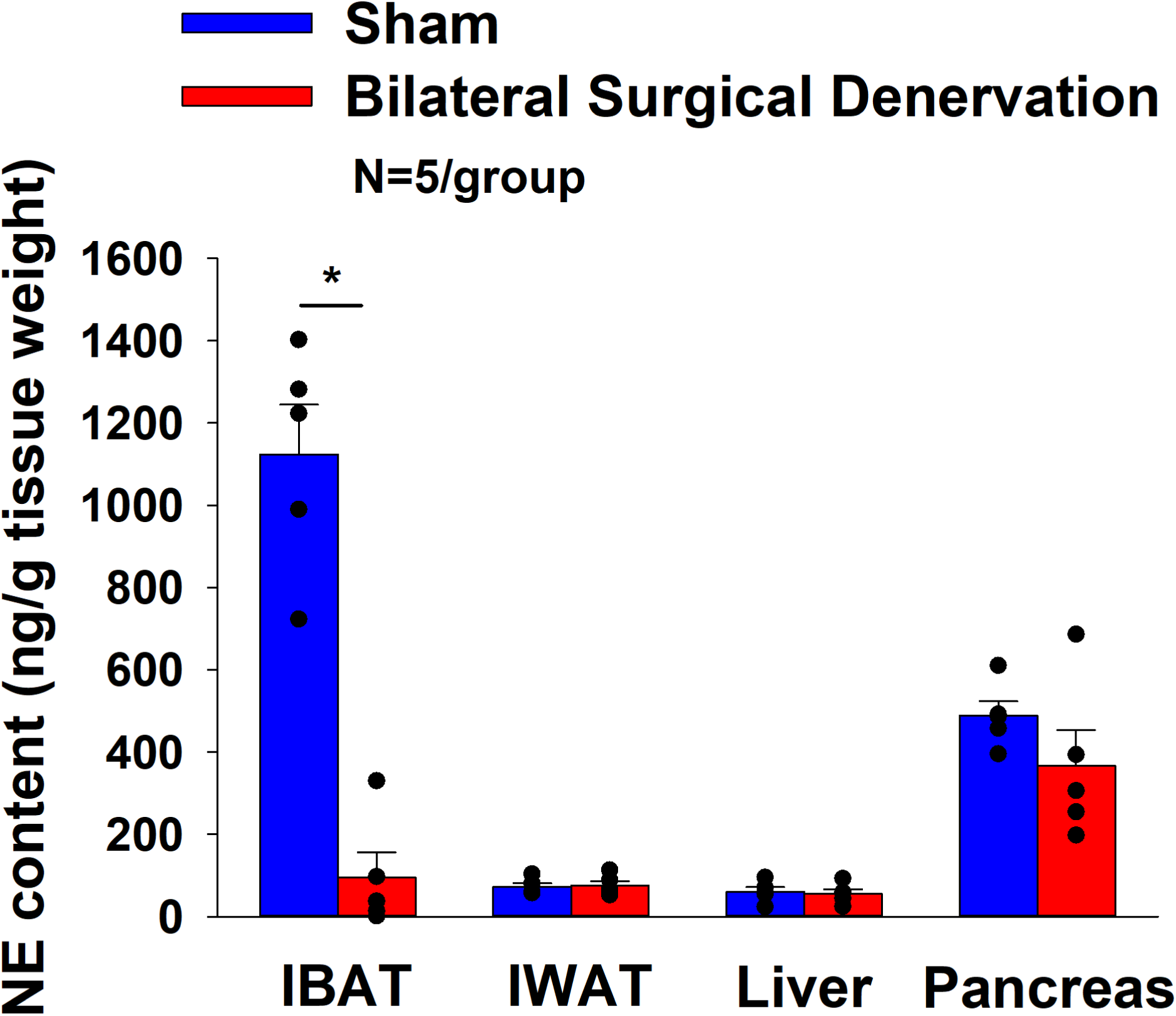
Effect of sham or IBAT surgical denervation procedure on IBAT NE content at 1-week post-sham or IBAT denervation in lean rats. Rats (N=10 total) were maintained on chow (13% kcal from fat) for approximately 1 week prior to undergoing a sham (N=5/group) or bilateral surgical IBAT denervation (N=5/group) procedures and were euthanized at 1-week post-sham or denervation procedure. NE content was measured from IBAT, IWAT and liver. Data are expressed as mean ± SEM. \**P*<0.05, †0.05<*P*<0.1 denervation vs sham.

### Study 2: Determine the success of the surgical denervation procedure at 1, 6 and 7-weeks post-sham or denervation in DIO rats by measuring NE content

Rats (N= 30 at study onset) were used for this study. The goal of this study was to 1) verify the success of the SNS denervation procedure in DIO rats by confirming a reduction of NE content that was specific to IBAT relative to other tissues (IWAT, liver and pancreas) and 2) confirm that these changes would persist for extended periods of time out to 7 weeks. By design, DIO rats were obese as determined by both BW (695 ± 13.9 g) and adiposity (230.1 ± 10.1 g fat mass; 32.7 ± 0.9% adiposity) after maintenance on the HFD for approximately 5 months prior to sham/denervation procedures. Sham and denervation groups were matched for BW, fat mass and lean mass such that there was no difference in baseline BW, lean mass, or fat mass between groups prior to surgery (*P*=NS).

All IBAT tissues from **Study 2** animals were analyzed for IBAT NE content and none of the fifteen animals were removed on account of having a failed IBAT denervation procedure. IBAT NE content was reduced by 94.7 ± 2.7, 96.8 ± 1.8 and 85.9±6.1% at 1-[(F(1,8) = 123.847, *P*=0.000)], 6-[(F(1,8) = 28.121, *P*=0.001)] and 7-weeks [(F(1,8) = 17.081, *P*<0.0003)] post-denervation (**Figure 2**) relative to IBAT NE content from sham operated rats. In contrast, NE content was unchanged in IWAT or liver in denervated rats relative to sham rats (*P*=NS) at 1, 6 or 7-weeks post-denervation. Similarly, there was also difference in NE content in pancreas at 6 or 7-weeks post-denervation relative to sham rats. However, there was a reduction of NE content in pancreas at 1-week post-denervation relative to sham rats (*P*<0.05).

**Figure 2:**
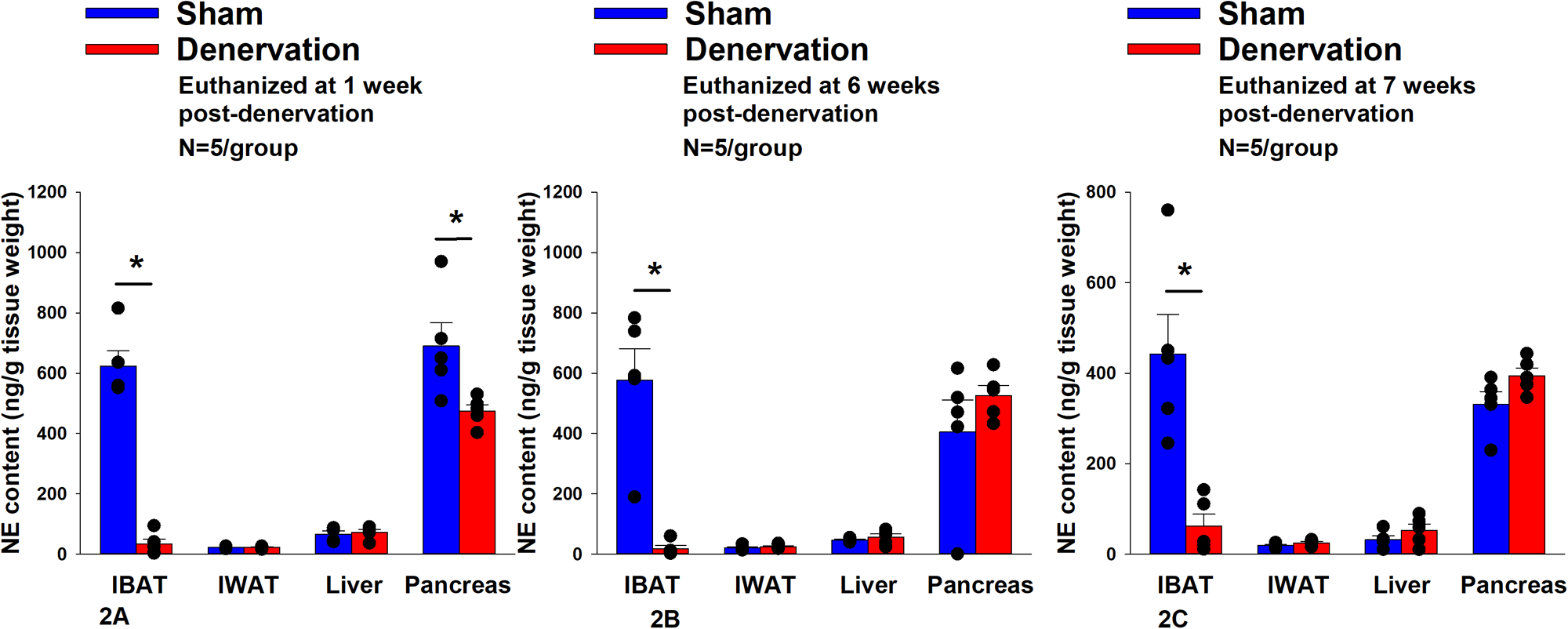
Effect of sham or IBAT surgical denervation procedure on IBAT NE content at 1-, 6- and 7-weeks post-sham or IBAT denervation in DIO rats. Rats (N=30 total) were maintained on a HFD (60% kcal from fat) for approximately 5 months prior to undergoing a sham (N=5/group/time) or bilateral surgical IBAT denervation (N=5/group/time) procedures and were euthanized at 1-, 6- and 7-weeks post-sham (N=15/group) or denervation procedure (N=15/group). NE content was measured from IBAT, IWAT, liver and pancreas. Data are expressed as mean ± SEM. \**P*<0.05, †0.05<*P*<0.1 denervation vs sham.

There were no significant differences in BWs between sham and denervation groups at 1, 6 or 7-week post-surgery (*P*=NS; data not shown). Similarly, there was no difference in fat mass, lean mass, percent fat mass or percent lean mass between groups at 1, 6 or 7-weeks post-denervation or sham surgeries (P=NS; data not shown). These findings are in agreement with others who have reported no difference in BW [70; 71; 72; 73; 74] or fat mass [71; 73] in hamsters or mice following bilateral surgical or chemical denervation of IBAT.

### Study 3: Determine if surgical denervation of IBAT changes the ability of the β3-AR agonist, CL 316243, to increase T_IBAT_ in DIO rats

Rats (N= 20 at study onset) were used for this study. The goal of this study was to confirm there was no functional defect in the ability of IBAT to respond to direct β3-AR stimulation because of the denervation procedure relative to sham operated animals. By design, DIO rats were obese as determined by both BW (684.1 ± 11.5 g) and adiposity (226.1 ± 8.1 g fat mass; 33 ± 0.8% adiposity) after maintenance on the HFD for approximately four months prior to sham/denervation procedures.

All IBAT tissues from **Study 3** animals were analyzed for IBAT NE content and 4 out of 10 animals were removed on account of having a failed IBAT denervation procedure. IBAT NE content was reduced in denervated rats by 87.5 ± 3.2% in denervated rats relative to sham-operated control rats [(F(1,12) = 67.209, *P*=0.000)]. In contrast, NE content was unchanged in IWAT, EWAT, liver or pancreas in denervated rats relative to sham rats (*P*=NS). There was no significant difference in BW between sham and denervation groups at the end of the study (≈ 4.5-weeks post-sham/denervation surgery) (*P*=NS; data not shown).

In sham rats, there was an overall effect of CL 316243 dose to increase T_IBAT_ when data were averaged over the 1-[(F(2,18) = 52.504, *P*<0.01] or 4-h post-injection period [(F(2,18) = 79.525, *P*<0.01]. Specifically, CL 316243 (1 mg/kg) increased T_IBAT_ at 0.25, 0.5, 0.75, 1, 1.25, 1.5, 1.75, 2, 3 and 4-h post-injection. The lowest dose (0.1 mg/kg) also stimulated T_IBAT_ at 0.25, 0.5, 0.75, 1, 1.25, 1.5, 1.75, 2, 3 and 4-h post-injection. CL 316243 also stimulated T_IBAT_ at 24-h post-injection at both doses (*P*<0.05; **Figure 3A**). Similar findings were apparent when measuring change in T_IBAT_ relative to baseline T_IBAT_ (**Figure 3B**).

**Figure 3A-F:**
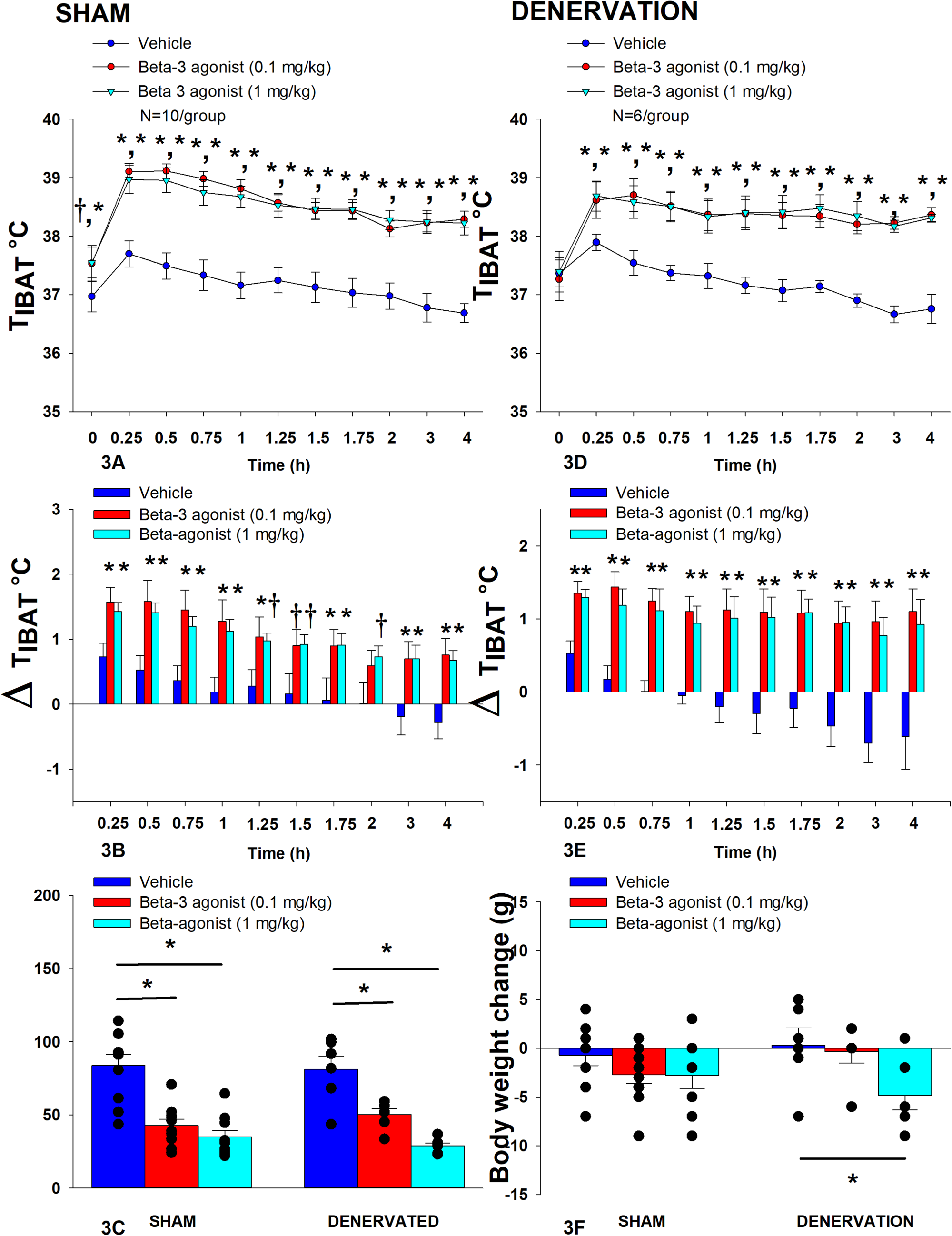
Effect of systemic β3-AR agonist (CL 316243) administration (0.1 and 1 mg/kg) on IBAT temperature (T_IBAT_), energy intake and BW post-sham or IBAT denervation in DIO rats. Rats (N=16 total) were maintained on HFD (60% kcal from fat; N=6–10/group) for approximately 4.25 months prior to undergoing a sham or bilateral surgical IBAT denervation and implantation of temperature transponders underneath IBAT. Animals were subsequently adapted to a 4-h fast prior to receiving IP injections of CL 316243 (0.1 or 1 mg/kg, IP) or vehicle (sterile water) where each animal received each treatment at approximately 7-day intervals. *A/C*, Effect of CL 316243 on T_IBAT_ in *A)* sham operated or *C)* IBAT denervated DIO rats; *B/D*, Effect of CL 316243 on change in T_IBAT_ relative to baseline T_IBAT_ (delta T_IBAT_) in *B)* sham operated or *D)* IBAT denervated DIO rats; *E*, Effect of CL 316243 on change in energy intake in sham or IBAT denervated DIO rats; *F*, Effect of CL 316243 on change in BW in sham or IBAT denervated DIO rats. Data are expressed as mean ± SEM. \**P*<0.05, †0.05<*P*<0.1 CL 316243 vs. vehicle.

Similarly, in denervated rats, there was an overall effect of CL 316243 dose to increase T_IBAT_ when data were averaged over the 1-[(F(2,10) = 17.714, *P*=0.001] or 4-h post-injection period [(F(2,10) = 36.703, *P*<0.01]. Specifically, CL 316243 (1 mg/kg) increased T_IBAT_ at 0.25, 0.5, 0.75, 1, 1.25, 1.5, 1.75, 2, 3 and 4-h post-injection. The lowest dose (0.1 mg/kg) also stimulated T_IBAT_ at 0.25, 0.5, 0.75, 1, 1.25, 1.5, 1.75, 2, 3 and 4-h post-injection. CL 316243 also stimulated T_IBAT_ at 24-h post-injection at the high dose (*P*<0.05; **Figure 3D**). Similar findings were apparent when measuring change in T_IBAT_ relative to baseline T_IBAT_ (**Figure 3E**).

Importantly, there was no difference in the T_IBAT_ response to CL 316243 (0.1 or 1 mg/kg) when the data were averaged over the 1-h or 4-h post-injection period between sham and denervated rats (*P*=NS). Overall, these findings indicate that IBAT denervation did not result in a change in the ability of CL 316243 to increase BAT thermogenesis (surrogate measure of EE) in DIO mice relative to sham operated rats.

#### Energy intake

In sham rats, CL 316243 reduced daily energy intake at both 0.1 and 1 mg/kg by 49 and 58.3% (P<0.05). Similarly, in denervated rats, CL 316243 also reduced daily energy intake at both 0.1 and 1 mg/kg (*P*<0.05) by 38 and 64.5% relative to vehicle (**Figure 3C**).

#### Body Weight (BW)

While CL 316243 did not impact BW in either group the high dose reduced BW gain in the denervated rats (*P*<0.05; **Figure 3F**).

Importantly, there was no difference in the effectiveness of CL 316243 (0.1 or 1 mg/kg) to reduce energy intake or weight gain between sham and denervated rats (*P*=NS). Overall, these findings indicate that IBAT denervation did not result in a significant change in the ability of CL 316243 to reduce energy intake or BW gain in DIO rats relative to sham operated rats.

### Study 4: Determine the extent to which OT-induced activation of sympathetic outflow to IBAT contributes to its ability to increase T_IBAT_ in DIO rats

Rats (N= 28 at study onset) were used for this study. After having confirmed there was no functional defect in the ability of IBAT to respond to direct β3-AR stimulation (**Study 3**), the goal of this study was to determine if OT-elicited elevation of T_IBAT_ requires intact SNS outflow to IBAT. DIO rats from **Study 3** were subsequently used in this study. Animals weighed approximately 631.6±10.3 g at the start of the studies. There was no significant difference in BW between sham and denervation groups at the end of the study (≍ 8.5-weeks post-sham/denervation surgery) (*P*=NS; data not shown).

In sham rats, there was an overall effect of 4V OT dose to increase T_IBAT_ when data were averaged over the 2-h post-injection period [(F(2,14) = 9.408, *P*=0.003]. Specifically, OT (5 μg) increased T_IBAT_ at 0.75, 1, 1.25, 1.5, and 1.75-h post-injection. The high dose produced a near significant stimulation of T_IBAT_ at 0.25 (0.05<*P*<0.1) and 2-h (*P*=0.052) post-injection. The lowest dose (1 μg) also stimulated T_IBAT_ at 1.75-h post-injection and produced a near significant stimulation of T_IBAT_ at 0.75-h post-injection (0.05<*P*<0.1; **Figure 4A**). Similar findings were apparent when measuring change in T_IBAT_ relative to baseline T_IBAT_ (**Figure 4B**).

**Figure 4A–D:**
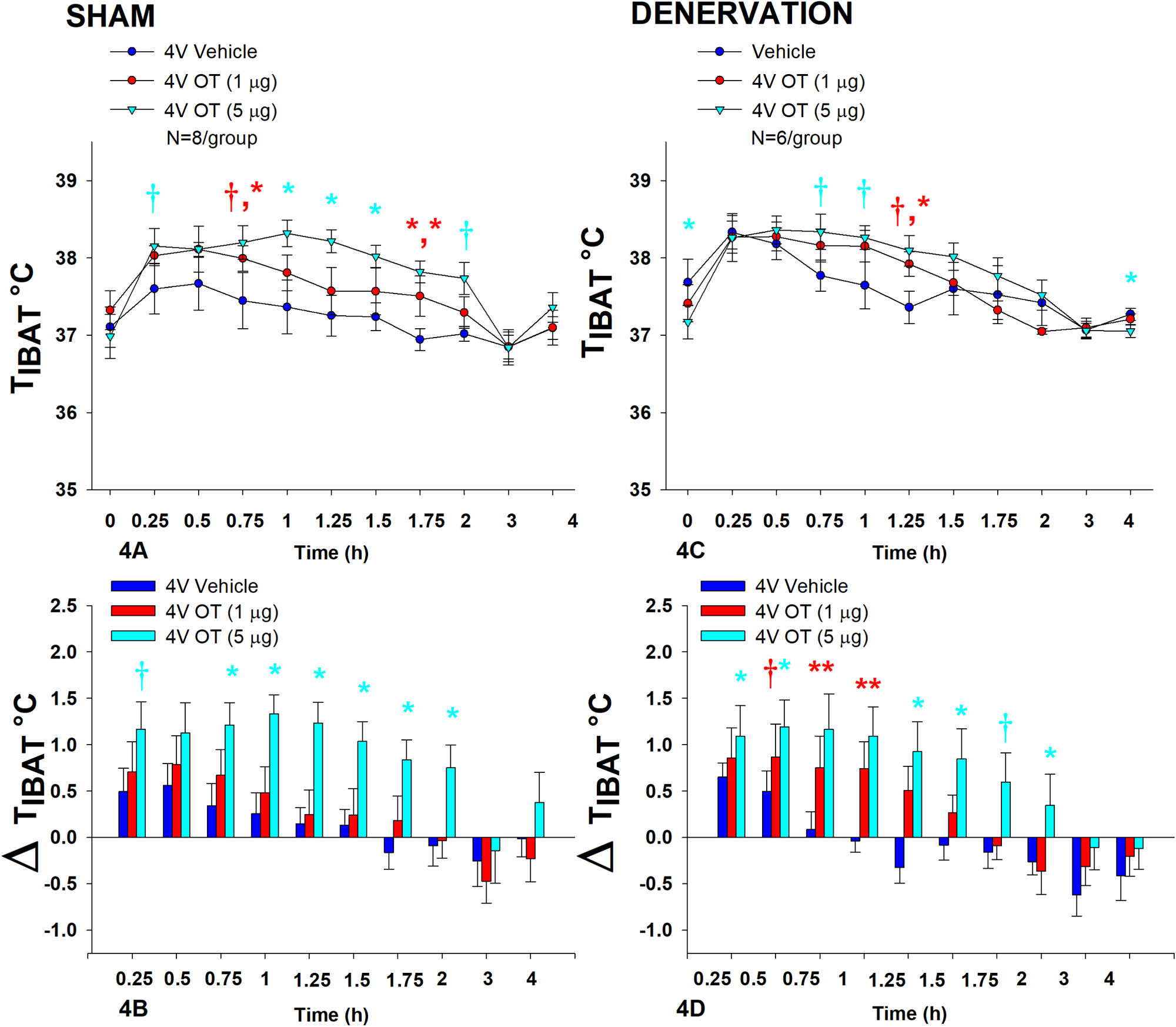
Effect of acute 4V OT administration (1 and 5 μg) on T_IBAT_ post-sham or IBAT denervation in male DIO rats. Rats (N=14 total) were maintained on HFD (60% kcal from fat; N=6–8/group) for approximately 4.25 months prior to undergoing a sham or bilateral surgical IBAT denervation and implantation of temperature transponders underneath IBAT. Rats were subsequently implanted with 4V cannulas and allowed to recover for 2 weeks prior to receiving acute 4V injections of OT or vehicle. Animals were subsequently adapted to a 4-h fast prior to receiving acute 4V injections of OT or vehicle *A/C*, Effect of acute 4V OT on T_IBAT_ in *A)* sham operated or *C)* IBAT denervated DIO rats; *B/D*, Effect of acute 4V OT on change in T_IBAT_ relative to baseline T_IBA_T (delta T_IBAT_) in *B)* sham operated or *D)* IBAT denervated DIO rats; Data are expressed as mean ± SEM. \**P*<0.05, †0.05<*P*<0.1 OT vs. vehicle.

In denervated rats, there was no overall effect of 4V OT dose to increase T_IBAT_ when data were averaged over the 2-h post-injection period [(F(2,10) = 1.174, *P*=0.348], likely due to the low dose (1 μg) not being able to produce a significant stimulation of T_IBAT_. Specifically, the high dose of OT (5 μg) increased T_IBAT_ at 1.25-h post-injection (*P*<0.05) and produced a near significant stimulation of T_IBAT_ at 0.75 and 1-h post-injection (0.05<*P*<0.1). The lowest dose (1 μg) produced a near significant stimulation of T_IBAT_ at 1.25-h post-injection (0.05<*P*<0.1; **Figure 4C**). Similar findings were apparent when measuring change in T_IBAT_ relative to baseline T_IBAT_ (**Figure 4D**) as well as when examining these effects in lean rats (**Supplemental Figure 1**).

Importantly, there was no difference in the T_IBAT_ response to 4V OT (5 μg) at 1.25-h post-injection between sham and denervated rats (*P*=NS). Overall, these findings indicate that IBAT denervation did not result in a significant change in the ability of 4V OT to increase BAT thermogenesis in denervated rats relative to sham operated rats.

### Study 5: Determine the extent to which OT-induced activation of sympathetic outflow to IBAT requires activation of β3-AR to increase T_IBAT_ in DIO rats

Rats (N= 8 at study onset) were used for this study. The goal of this study was to determine whether hindbrain administration of OT required activation of β3-AR to stimulate BAT thermogenesis. We initially identified a dose of the β3-AR antagonist, SR 59230A, that was sufficient to block the effects of the β3-AR agonist, CL316243, to stimulate T_IBAT_ at 0.5-h and 0.75-h post-CL316243 treatment (P=NS vehicle vs CL316243; **Supplemental Figure 2**). This dose was then used to address whether 4V OT stimulated T_IBAT_ through a β3-AR driven mechanism.

4V OT (in the absence of the β3-AR antagonist) increased T_IBAT_ at 0.75, 1, 1.25, 1.75, 2, and 4-h post-injection (*P*<0.05) and produced a near significant stimulation of T_IBAT_ at 1.5, and 1.25-h post-injection (0.05<*P*<0.1). Following β3-AR antagonist pre-treatment, 4V OT stimulated T_IBAT_ at 0.75 (*P*=0.029), 1 (*P*=0.022), 1.25 (*P*=0.005), 1.5-h (*P*=0.003), and 1.75 (*P*=0.009) post-injection and produced a near significant stimulation of T_IBAT_ at 0.25 (*P*=0.064) post-injection (**Figure 5**). 4V OT also appeared to stimulate T_IBAT_ at 0.5 (*P*=0.109) and 2-h post-injection (*P*=0.125).

**Figure 5A–B:**
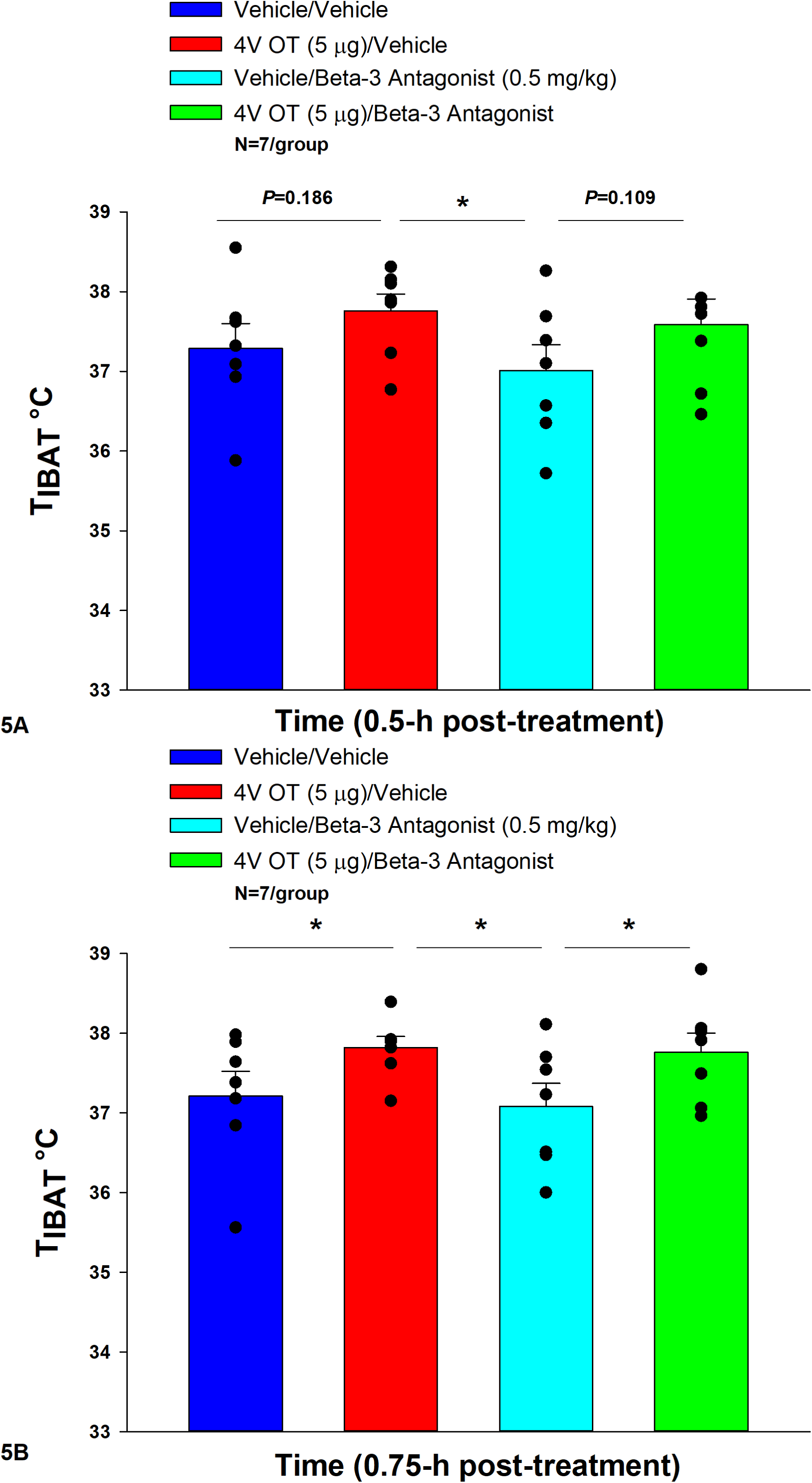
Effect of β3-AR antagonism on ability of acute 4V OT to increase T_IBAT_ in DIO rats. Rats (N=7 total) were maintained on HFD (60% kcal from fat; N=7/group) for approximately 4 months prior to being implanted with temperature transponders underneath the left IBAT depot. Rats were subsequently implanted with 4V cannulas and allowed to recover for 4 weeks. Animals were subsequently adapted to a 4-h fast and handling during the week prior to the experiment. On an experimental day, 4-h fasted rats received the β3-AR antagonist, SR 59230A (0.5 mg/kg, IP) or vehicle (DMSO) approximately 20 min prior to 4V administration of either OT (5 μg/μl) or vehicle. *A*, Effect of systemic pre-treatment with the β3-AR antagonist, SR 59230A, on the ability of acute 4V OT to stimulate T_IBAT_ at 0.5-h post-injection in DIO rats; B), Effect of systemic pre-treatment with the β3-AR antagonist, SR 59230A, on the ability of acute 4V OT to stimulate T_IBAT_ at 0.75-h post-injection in DIO rats. Data are expressed as mean ± SEM. \**P*<0.05, †0.05<*P*<0.1.

Two-way repeated-measures ANOVA revealed a significant main effect of OT to increase T_IBAT_ at 0.75-h post-injection [F(1,18) = 10.111, *P*=0.005]. There was no significant main effect of the β3-AR antagonist on T_IBAT_ at 0.75-h post-injection [F(1,18) = 0.221, *P*=0.644]. There was also no significant interaction between OT and the β3-AR antagonist on T_IBAT_ at 0.75-h post-injection [F(1,18) = 0.030, *P*=0.864].

We also found that 4V OT (5 μg) was effective (n=11; data not shown) at stimulating T_IBAT_ in lean rats even in the presence of the β3-AR antagonist, SR 59230A, that was given at 2-(1 mg/kg) and 10-fold higher (5 mg/kg) doses approximately 12 minutes prior to 4V OT (n=8; data not shown). Note that in contrast to the study design in the DIO rats, the timing of the drug treatments in the lean rats occurred toward the end of the light cycle.

Overall, these findings suggest that OT does not require activation of β3-AR to stimulate BAT thermogenesis.

### Study 6A: Determine the extent to which OT-induced activation of sympathetic outflow to IBAT contributes to its ability to impact BW in DIO rats

Rats (N= 43 at study onset) were used for this study. The goal of this study was to determine if OT-elicited weight loss requires intact SNS outflow to IBAT. By design, DIO rats were obese as determined by both BW (682.2 ± 9.7 g) and adiposity (211.8 ± 6.8 g fat mass; 30.9 ± 0.7% adiposity) after maintenance on the HFD for at least 4.5 months prior to sham/denervation procedures. One subset (N=4) was maintained on the HFD for approximately 6 months prior to sham/denervation procedures and animals were divided equally among treatment groups. All IBAT tissues from **Study 6A** animals were analyzed for IBAT NE content and 3 of the 19 IBAT denervated animals were removed on account of having a failed IBAT denervation procedure. IBAT NE content was reduced in denervated rats by 91.1 ± 2.4% in denervated rats relative to sham-operated control rats [(F(1,38) = 81.864, *P*<0.01)]. In contrast, NE content was unchanged in EWAT, liver or pancreas in denervated rats relative to sham rats (*P*=NS). In contrast, there was a near significant increase (≈1.35-fold) in NE content in IWAT in denervated rats relative to sham rats [(F(1,38) = 4.116, *P*=0.05)]. There was no significant difference in BW between sham and denervation groups at the end of the study (≈ 7-8-weeks post-sham/denervation surgery) (*P*=NS; data not shown).

As expected, in sham rats, 4V vehicle resulted in 6.6 ± 2.1% weight gain relative to vehicle pre-treatment [(F(1,9) = 9.610, *P*=0.013)]. In contrast, 4V OT failed to reduce BW relative to OT pre-treatment [(F(1,9) = 0.582, *P*=0.465)] (**Figure 6A**) but it reduced weight gain (**Figure 6B**) throughout the 29-day infusion period. OT treatment reduced weight gain on day 3 and 7-29 (*P*<0.05) and produced a near significant reduction of weight gain on day 6 (*P*=0.088). By the end of the infusion period (infusion day 29), OT had reduced weight gain by -7.2 ± 9.6 g relative to vehicle treated animals 49.1 ± 10.0 g (*P*<0.05). OT reduced relative fat mass (pre- vs post-intervention) (**Figure 6C**; *P*<0.05), fat mass and relative lean mass (pre- vs post-intervention) but had no effect on total lean body mass (*P*=NS). These effects that were mediated, at least in part, by a modest reduction of energy intake that was evident during weeks 2 and 4 of the treatment period (**Figure 6D**; *P*<0.05). OT also produced a near significant reduction of energy intake at week 3 of the treatment period (*P*=0.062).

**Figure 6A–D:**
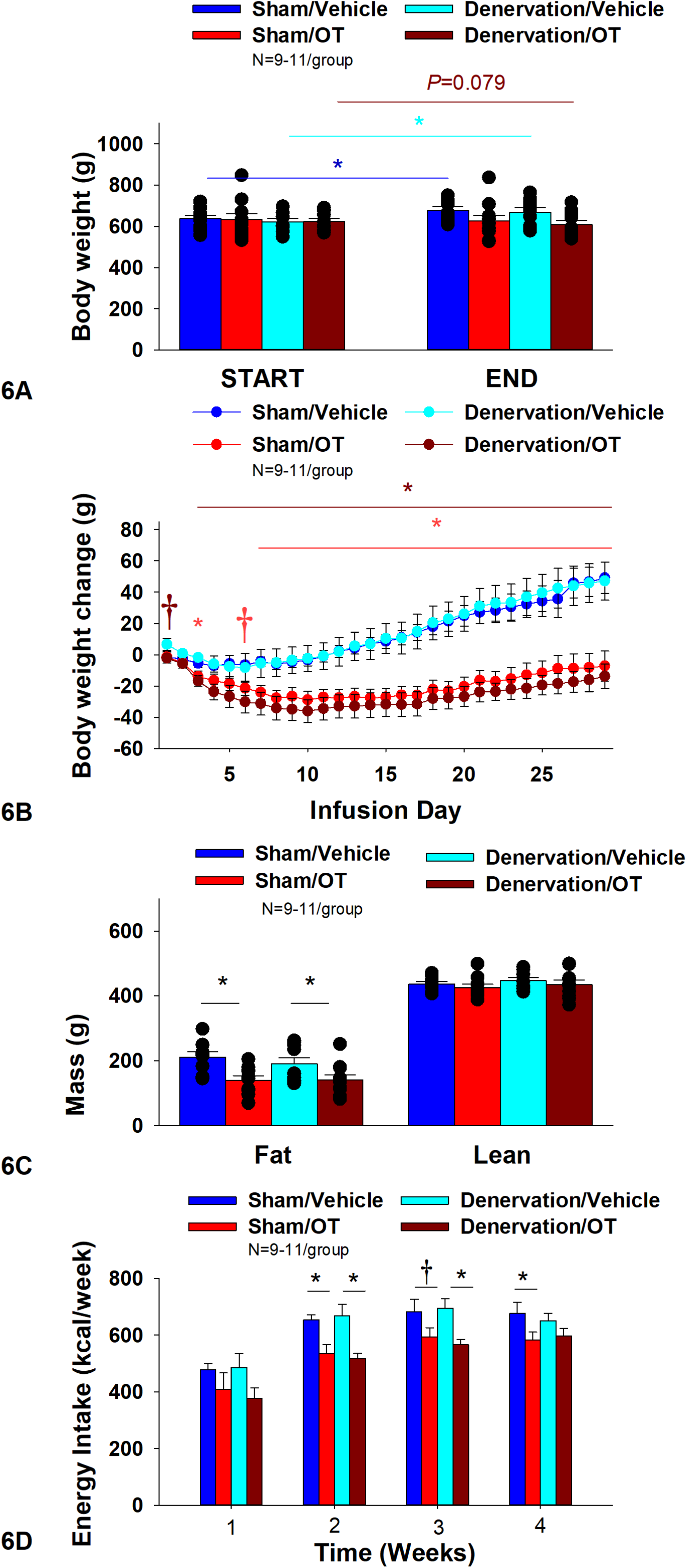
Effect of chronic 4V OT infusions (16 nmol/day) on BW, adiposity and energy intake post-sham or IBAT denervation in male DIO rats. *A*, Rats (N=38 total) were maintained on HFD (60% kcal from fat; N=8–11/group) for approximately 4.5 months prior to undergoing a sham or bilateral surgical IBAT denervation. Rats were subsequently implanted with 4V cannulas and allowed to recover for 2 weeks prior to being implanted with subcutaneous minipumps that were subsequently attached to the 4V cannula. *A*, Effect of chronic 4V OT or vehicle on BW in sham operated or IBAT denervated DIO rats; *B*, Effect of chronic 4V OT or vehicle on BW change in sham operated or IBAT denervated DIO rats; *C*, Effect of chronic 4V OT or vehicle on adiposity in sham operated or IBAT denervated DIO rats; *D*, Effect of chronic 4V OT or vehicle on adiposity in sham operated or IBAT denervated DIO rats. Data are expressed as mean ± SEM. \**P*<0.05, †0.05<*P*<0.1 OT vs. vehicle.

In denervated rats, 4V vehicle resulted in an expected 7.6 ± 2.0% weight gain relative to vehicle pre-treatment [(F(1,8) = 12.617, *P*=0.007)]. In contrast, 4V OT produced a near significant reduction of BW by 2.9 ± 1.4% relative to OT pre-treatment [(F(1,9) = 3.923, *P*=0.079)]. (**Figure 56**) and reduced weight gain (**Figure 6B**) throughout the 29-day infusion period. OT treatment reduced weight gain on days 3–29 (*P*<0.05). By the end of the infusion period (infusion day 29), OT had reduced weight gain by –13.8 ± 7.9 g relative to vehicle treated animals 47.1 ± 12.3 g (*P*<0.05). OT reduced relative fat mass (pre- vs post-intervention) (**Figure 6C**; *P*<0.05) and fat mass but had no effect on relative or total lean body mass (*P*=NS). These effects that were mediated, at least in part, by a modest reduction of energy intake that persisted during weeks 2 and 3 of the treatment period (**Figure 6D**; *P*<0.05). Similar effects were also observed in rats with more pronounced obesity (**Supplemental Figure 3**).

Two-way ANOVA revealed a significant main effect of OT to reduce body weight gain on day 29 [F(1,33) = 34.603, *P*<0.01] but no overall effect of denervation [F(1,33) = 0.185, *P*=0.670] or an interactive effect between OT and denervation on body weight gain on day 29 [F(1,33) = 0.053, *P*=0.820]. Similarly, two-way ANOVA revealed a significant main effect of OT to reduce fat mass [F(1,34) = 13.931, *P*<0.01] but no overall effect of denervation [F(1,34) = 0.405, *P*=0.529] or an interactive effect between OT and denervation on fat mass [F(1,34) = 0.470, *P*=0.498]. In addition, we found a significant main effect of OT to energy intake (week 2) [F(1,36) = 21.530, *P*<0.01] but no overall effect of denervation [F(1,36) = 0.004, *P*=0.948] or an interactive effect between OT and denervation on energy intake (week 2) [F(1,36) = 0.295, *P*=0.590].

Importantly, there was no difference in the effectiveness of 4V OT to reduce weight gain at the end of the infusion period (day 29), energy intake, and relative fat mass or fat mass between sham and denervated rats (*P*=NS). Based on these collective findings, we conclude that SNS innervation of IBAT is not a predominant contributor of OT-elicited reduction of weight gain and adiposity.

#### Adipocyte size

There was a near significant effect of denervation to reduce adipocyte size in IWAT (*P*=0.053) but no effect of OT to significantly impact IWAT adipocyte size in either the sham or denervation group (*P*=NS) (**Figure 7A**). There was a significant effect of denervation to reduce adipocyte size in EWAT (*P*=0.049; **Figure 7B**). In addition, there was a near significant effect of 4V OT to reduce adipocyte size in EWAT in both sham (*P*=0.133) and denervated rats (*P*=0.104) (**Figure 7B**).

**Figure 7A–B:**
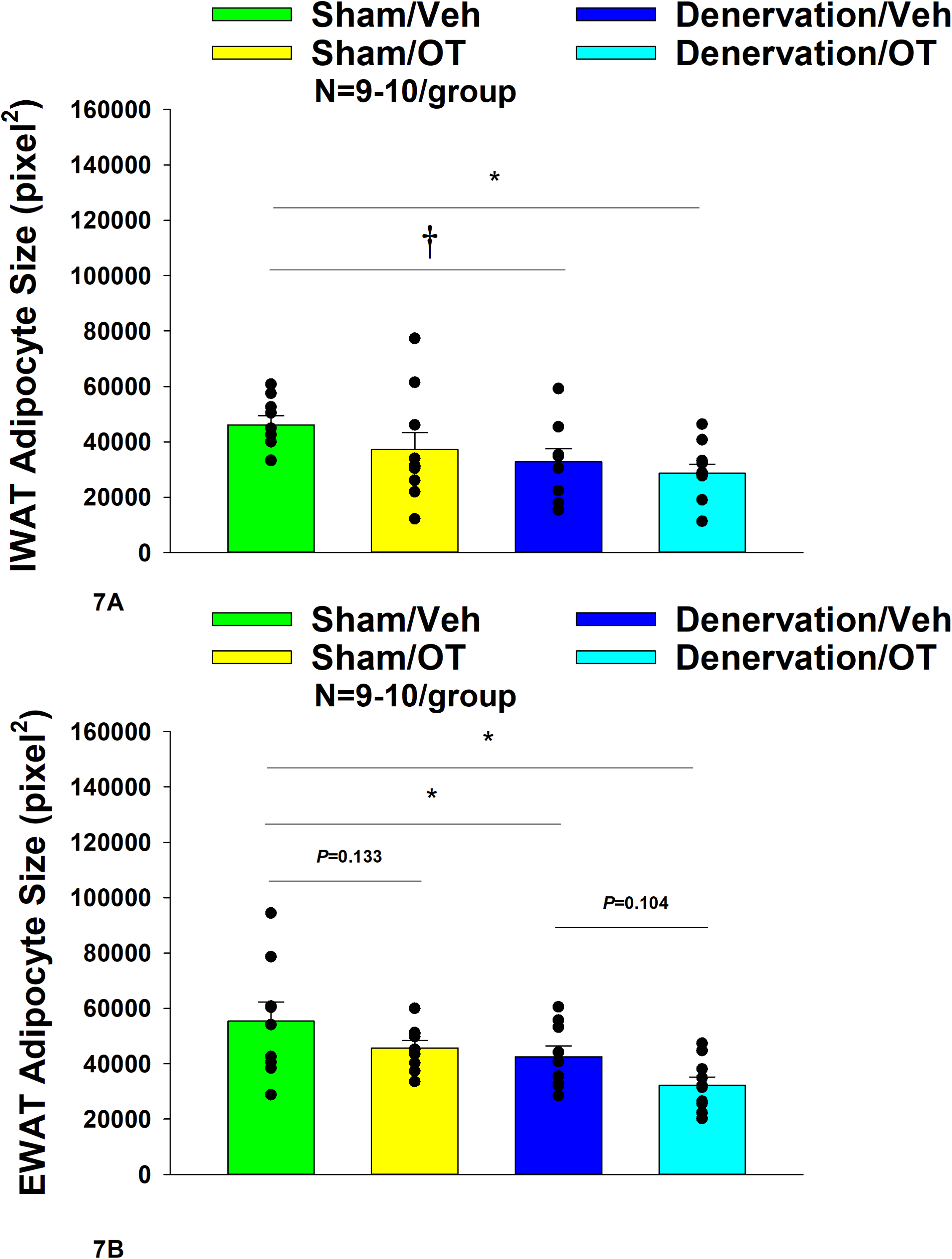
Effect of chronic 4V OT infusions (16 nmol/day) on adipocyte size post-sham or IBAT denervation in male DIO rats. *A*, Adipocyte size (pixel^2^) was measured in IWAT from rats (N=36 total) that received chronic 4V infusion of OT (16 nmol/day) or vehicle in sham or IBAT denervated DIO rats (N=4–5/group). *B*, Adipocyte size was measured in EWAT from rats (N=35 total) that received chronic 4V infusion of OT (16 nmol/day) or vehicle in sham operated or IBAT denervated rats (N=4–5/group). Data are expressed as mean ± SEM. \**P*<0.05 OT vs. vehicle.

#### Plasma hormone concentrations

To characterize the endocrine and metabolic effects between sham and denervated DIO rats, we measured blood glucose levels and plasma concentrations of leptin, insulin, FGF-21, irisin, adiponectin, TC, and FFAs at various times post-sham and denervation procedure. At 1-week post-denervation/sham, there was a significant reduction of blood glucose in the denervation group relative to the sham group (*P*<0.05). In contrast, there were no differences in any of the other metabolic measures between respective sham and denervated animals at 1, 6 or 7-weeks post-sham or denervation surgery (**Table 1**).

**Table 1.** Changes in plasma hormones at 1, 6, and 7-weeks post-sham or IBAT denervation in DIO rats. Blood was collected by tail vein (blood glucose) and cardiac stick (plasma hormones) following a 4-h fast. Different letters denote significant differences between respective sham and denervation groups at 1, 6 or 7-weeks post-sham or denervation. Shared letters denote no significant differences between respective sham and denervation groups. Data are expressed as mean ± SEM (N=5/group; N=28-30 total).

|  | 1 week |  | 6 weeks |  | 7 weeks |  |
| --- | --- | --- | --- | --- | --- | --- |
|  | Sham | Denervation | Sham | Denervation | Sham | Denervation |
| Leptin (ng/mL) | 25.7 ± 4.4<sup>a</sup> | 31.9 ± 5.8<sup>a</sup> | 59.7 ± 10.8<sup>a</sup> | 53.4 ± 10.0<sup>a</sup> | 50.2 ± 5.3<sup>a</sup> | 55.3 ± 9.3<sup>a</sup> |
| Insulin (ng/mL) | 3.4 ± 0.6<sup>a</sup> | 5.0 ± 1.8<sup>a</sup> | 5.6 ± 0.7<sup>a</sup> | 9.1 ± 3.8<sup>a</sup> | 6.0 ± 0.7<sup>a</sup> | 9.5 ± 2.3<sup>a</sup> |
| FGF-21 (pg/mL) | 181.5 ± 20.4<sup>a</sup> | 211.7 ± 68.1<sup>a</sup> | 219.3 ± 39.3<sup>a</sup> | 180.9 ± 53.2<sup>a</sup> | 212 ± 63.2<sup>a</sup> | 240.5 ± 86.2<sup>a</sup> |
| Irisin (μg/mL) | 11.4 ± 2.6<sup>a</sup> | 10.3 ± 0.8<sup>a</sup> | 10.6 ± 1.2<sup>a</sup> | 11.3 ± 0.8<sup>a</sup> | 10.6 ± 0.3<sup>a</sup> | 7.7 ± 0.9<sup>a</sup> |
| Adiponectin (μg/mL) | 7.4 ± 1.0<sup>a</sup> | 7.1 ± 0.4<sup>a</sup> | 8.1 ± 0.5<sup>a</sup> | 7.5 ± 0.5<sup>a</sup> | 7.2 ± 0.7<sup>a</sup> | 7.3 ± 0.8<sup>a</sup> |
| Blood Glucose (mg/dL) | 169.8 ± 11.9<sup>a</sup> | 140.4 ± 6.2<sup>b</sup> | 152 ± 8.2<sup>a</sup> | 169.6 ± 7.2<sup>a</sup> | 156 ± 11.4<sup>a</sup> | 161.2 ± 5.1<sup>a</sup> |
| TG | 78.5 ± 11.4<sup>a</sup> | 97.3 ± 13.1<sup>a</sup> | 119.9 ± 11.6<sup>a</sup> | 150.9 ± 40.2<sup>a</sup> | 101.9 ± 12.4<sup>a</sup> | 130.7 ± 8.6<sup>a</sup> |
| FFA (mEq/L) | 0.19 ± 0.01<sup>a</sup> | 0.23 ± 0.03<sup>a</sup> | 0.22 ± 0.02<sup>a</sup> | 0.25 ± 0.03<sup>a</sup> | 0.27 ± 0.06<sup>a</sup> | 0.22 ± 0.04<sup>a</sup> |
| Total Cholesterol (mg/dL) | 80.1 ± 3.1<sup>a</sup> | 96.8 ± 9.3<sup>a</sup> | 137.3 ± 8.7<sup>a</sup> | 126.5 ± 7.3<sup>a</sup> | 126.4 ± 15.5<sup>a</sup> | 130.2 ± 9.3<sup>a</sup> |

We also characterized the endocrine and metabolic effects of 4V OT (16 nmol/day) in sham and denervated DIO rats. We found that denervation alone in vehicle-treated animals resulted in a near significant reduction of plasma irisin relative to sham control animals (*P*=0.056). In addition, we found OT treatment was associated with a reduction of plasma leptin in the sham group (*P*<0.05) which coincided with OT-elicited reductions in fat mass. In addition, OT treatment was associated with a near significant reduction of plasma adiponectin in the denervation group (*P*=0.057) (**Table 2**).

**Table 2.** Plasma measurements following 4V infusions of OT (16 nmol/day) or vehicle in sham and IBAT denervated DIO rats. Blood was collected by tail vein (blood glucose) and cardiac stick (plasma hormones) following a 4-h fast. Different letters denote significant differences between treatments. Shared letters denote no significant differences between treatments. Data are expressed as mean ± SEM (N=8-11/group; N=38 total).

| **4V** | **VEH** | **OT** | **VEH** | **OT** |
| --- | --- | --- | --- | --- |
|  | **Sham** | **Sham** | **Denervation** | **Denervation** |
| **Leptin (ng/mL)** | 44.9 ± 4.8<sup>a</sup> | 29 ± 3.2<sup>b</sup> | 36.8 ± 6.2<sup>ab</sup> | 28.2 ± 5.9<sup>b</sup> |
| **Insulin (ng/mL)** | 6.0 ± 1.0<sup>a</sup> | 5.0 ± 0.6<sup>a</sup> | 4.9 ± 0.7<sup>a</sup> | 3.9 ± 0.7<sup>a</sup> |
| **FGF-21 (pg/mL)** | 153.2 ± 29.3<sup>a</sup> | 166.2 ± 40.7<sup>a</sup> | 140.1 ± 25.2<sup>a</sup> | 130.9 ± 19.7<sup>a</sup> |
| **Irisin (µg/mL)** | 8.2 ± 0.8<sup>ab</sup> | 8.5 ± 0.6<sup>a</sup> | 6.2 ± 0.4<sup>b</sup> | 6.6 ± 0.8<sup>ab</sup> |
| **Adiponectin (µg/mL)** | 7.4 ± 0.6<sup>a</sup> | 6.6 ± 0.4<sup>a</sup> | 7.5 ± 0.3<sup>a</sup> | 6.1 ± 0.6<sup>b</sup> |
| **Blood Glucose (mg/dL)** | 183.5 ± 7.3<sup>a</sup> | 188.4 ± 4.4<sup>a</sup> | 184.0 ± 8.3<sup>a</sup> | 178.8 ± 7.1<sup>a</sup> |
| **FFA (mEq/L)** | 0.24 ± 0.03<sup>a</sup> | 0.24 ± 0.02<sup>a</sup> | 0.28 ± 0.04<sup>a</sup> | 0.22 ± 0.03<sup>a</sup> |
| **Total Cholesterol (mg/dL)** | 116.4 ± 7.1<sup>a</sup> | 113.1 ± 3.3<sup>a</sup> | 106.9 ± 9.4<sup>a</sup> | 101.1 ± 6.5<sup>a</sup> |

### Study 6B: Determine the success of the surgical denervation procedure at 6-8 weeks post-sham or denervation in DIO rats by measuring thermogenic gene expression

The goal of this study was to 1) confirm the success of the SNS denervation of IBAT procedure in DIO rats using an independent and complementary marker to IBAT NE content (IBAT thermogenic gene expression), 2) verify that these changes would persist out to more prolonged periods of time (6–8 weeks) and 3) determine if changes in thermogenic gene expression in EWAT occur in response to IBAT denervation.

#### IBAT

There was a significant reduction of IBAT Dio2 mRNA expression (*P*=0.016) in denervated rats relative to IBAT from sham operated rats (*P*<0.05; **Table 3**). In addition, there was a near significant reduction of IBAT UCP-1 (*P*=0.057) and Prdm16 mRNA (*P*=0.052) expression from denervated rats relative to sham operated rats (**Table 3**).

**Table 3.**
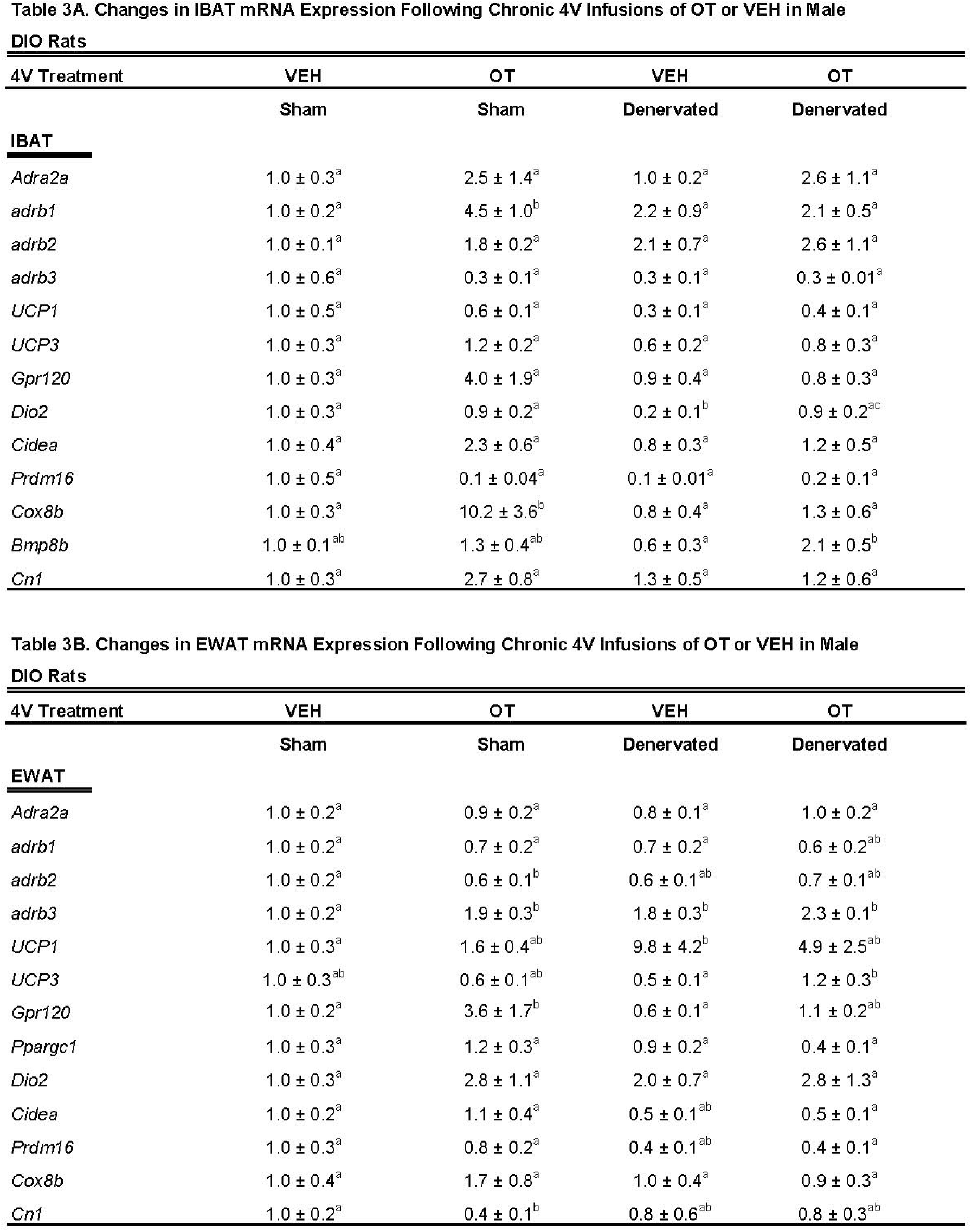
Changes in IBAT and EWAT gene expression following 4V infusions of OT or vehicle in male sham or IBAT denervated DIO rats. IBAT and EWAT were collected following a 4-h fast. *A*, Changes in IBAT mRNA expression following 4V infusions of OT or vehicle in male sham or IBAT denervated DIO rats (N=13-37 total); *B*, Changes in EWAT mRNA expression following 4V infusions of OT or vehicle in male sham or IBAT denervated DIO rats (N=23-29 total). Different letters denote significant differences between treatments. Shared letters denote no significant differences between treatments. Data are expressed as mean ± SEM (N=3-10/group).

#### EWAT

There was an increase of EWAT UCP-1 (*P*=0.024) and β3-AR (*P*=0.032) mRNA expression in denervated rats compared to sham rats. In addition, there was a near significant reduction of IBAT β2-AR (*P*=0.073) and Prdm16 (*P*=0.078) mRNA expression in denervated rats compared to sham operated rats (**Table 3**).

Collectively, these findings indicate that IBAT denervation results in significant (*Dio2*) or near significant reductions (*Ucp1* and *Prdm16*) of thermogenic gene markers in IBAT similar to what we recently reported following IBAT denervation in mice [29]. In addition, these findings indicate that IBAT denervation results in significant increases in the thermogenic markers, *Ucp1* and *Adrb3* in EWAT (indicative of browning of WAT).

### Study 6C: Determine the extent to which IBAT denervation contributes to the ability of OT to impact thermogenic gene expression in IBAT and EWAT in DIO rats

The goal of this study was to determine 1) if 4V OT elicits changes in thermogenic gene expression in IBAT and EWAT in sham operated rats, and 2) if 4V OT elicits changes in thermogenic gene expression in IBAT and EWAT in IBAT denervated rats.

#### IBAT

We found that chronic 4V OT treatment elicited a significant increase in the expression of IBAT Cox8b (*P*=0.004) and β1-AR mRNA expression (*P*=0.004) but this effect was blocked in denervated rats. 4V OT also elicited a near significant increase in IBAT Gpr120 (*P*=0.063), Cidea (*P*=0.075) and CN1 (*P*=0.069) mRNA expression and a reduction of IBAT Prdm16 mRNA expression (*P*=0.068) in sham operated rats. There was also a significant effect of 4V OT to increase IBAT Dio2 (*P*=0.041) and Bmp8b (*P*=0.019) mRNA in denervated rats.

In addition, 4V OT produced significant (*Cox8b*, *Adrb1*) or near significant increases (*Gpr120*, *Cidea* and *CN1*) in thermogenic markers in IBAT in sham rats, some of which were blocked in response to denervation of IBAT (*Cox8b*, *Adrb1*). In contrast, 4V OT increased mRNA expression of Dio2 and Bmp8b in denervated rats but not in sham rats raising the possibility that activation of these thermogenic markers might contribute to the effects of 4V OT on BAT thermogenesis in denervated animals.

#### EWAT

4V OT elicited a significant increase in EWAT β3-AR (*P*=0.018) and Gpr120 (*P*=0.03) mRNA expression in sham rats while it also reduced EWAT CN1 mRNA expression (*P*=0.051) in sham rats. In addition, 4V OT also increased EWAT UCP-3 (*P*=0.042) mRNA expression in denervated rats.

4V OT resulted in a significant increase (*Adrb3*, *Gpr120*) or decrease (*CN1*) of thermogenic markers in EWAT of sham operated rats while it also increased the thermogenic marker (*Ucp3*) in denervated rats. These findings raise the possibility that different thermogenic markers in EWAT may contribute, in part, to the metabolic effects of 4V OT in sham (*Adrb3*, *Gpr120*, *CN1*) and denervated rats (*Ucp3*).

### Study 7A: Determine the extent to which 4V OT increases gross motor activity, core temperature and T_IBAT_ in DIO rats

Rats (N= 19 at study onset) were used for this study. The goal of this study was to determine if hindbrain administration of OT increases gross motor activity at doses that also increase BAT thermogenesis.

#### Gross motor activity

Using repeated measures ANOVA, we found a near significant overall effect of 4V OT dose to stimulate gross motor activity [(F(18,162) = 1.430, *P* = 0.124] across ten time points over the 4-h measurement period. Specifically, 4V OT (5 μg/μL) increased activity at 1.5-h post-injection (*P*<0.05) and produced a near significant stimulation of activity at 0.5, 0.75-h post-injection (0.05<*P*<0.1) (**Figure 8A**). The lower dose of OT (1 μg/μL) also produced a near significant increase of activity at 0.75-h post-injection (0.05<*P*<0.1). 4V OT (5 μg/μL) also reduced activity at 4-h post-injection. However, 4V OT (5 μg/μL) also appeared to elicit a modest increase in gross motor activity when the data were averaged over the 1 (*P*=0.118) or 2-h (*P*=0.172) post-injection period but these effects were not significant (data not shown).

**Figure 8A–D:**
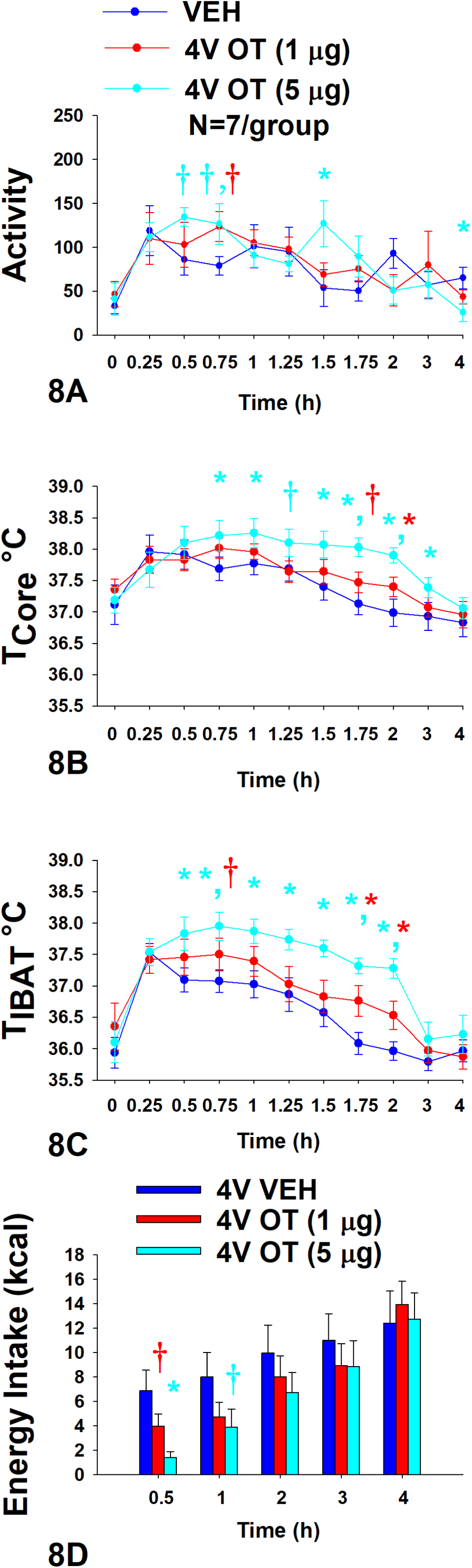
Effect of acute 4V OT on gross motor activity, core temperature, T_IBAT_ and energy intake in DIO rats. Rats (N=7 total) were maintained on HFD (60% kcal from fat; N=7/group) for approximately 6 months prior to being implanted with temperature transponders underneath IBAT, intra-abdominal telemetry devices and 4V cannulas. Animals were subsequently adapted to a 4-h fast prior to receiving acute 4V injections of OT or vehicle. Animals remained fasted for additional 4 h during which time gross motor activity, T_IBAT_ and core temperature was measured. Energy intake data was collected separately from the same animals following a 4-h fast *A*, Effect of acute 4V OT on gross motor activity in DIO rats; *B*, Effect of acute 4V OT on T_IBAT_ in DIO rats; C, Effects of acute 4V OT on core temperature in DIO rats, and D, Effect of acute 4V OT on energy intake in DIO rats. Data are expressed as mean ± SEM. \**P*<0.05, †0.05<*P*<0.1 OT vs. vehicle.

#### Core temperature

Using repeated measures ANOVA, we also found an overall effect of 4V OT dose to stimulate core temperature [(F(18,162) = 3.203, *P* < 0.01] across ten time points over the 4-h measurement period. Specifically, 4V OT (5 μg/μL) increased core temperature at 0.75, 1, 1.5, 1.75, 2, and 3-h post-injection (*P*<0.05) (**Figure 8B**) and produced a near significant increase of core temperature at 1.25-h post-injection (0.05<*P*<0.1). The lower dose of OT (1 μg/μL) also increased core temperature at 2-h post-injection (*P*<0.05) and produced a near significant increase of core temperature at 1.75-h post-injection (0.05<*P*<0.1). While there was no overall effect of 4V dose to increase core temperature when averaged over 1-h post-injection period (CT_1-hour AVE_), there was a significant overall effect of 4V OT dose to increase core temperature when the data were averaged over the 2 (CT_2-hour AVE_) [(F(2,12) = 6.584, *P*=0.02], 3 [(F(2,12) = 4.556, *P*=0.034] (CT_3-hour AVE_) and 4-h (CT_4-hour AVE_) [(F(2,12) = 5.523, *P*=0.020] post-injection (data not shown). Specifically, the high dose of OT (5 μg/μL) was able to stimulate CT_2-hour AVE_, CT_3-hour AVE_, and CT_4-hour AVE_ (*P*<0.05) while the lower dose was ineffective (data not shown; *P*=NS).

#### T_IBAT_

Using repeated measures ANOVA, we also found an overall effect of 4V OT dose to stimulate T_IBAT_ [(F(18,162) = 1.917, *P* = 0.018] across ten time points over the 4-h measurement period. Specifically, 4V OT (5 μg/μL) increased T_IBAT_ at 0.5, 0.75, 1, 1.25, 1.5, 1.75, and 2-h post-injection (*P*<0.05) (**Figure 8C**). The lower dose of OT (1 μg/μL) also increased T_IBAT_ at 1.75 and 2-h post-injection (*P*<0.05) and produced a near significant increase of T_IBAT_ at 0.75-h post-injection (0.05<*P*<0.1). There also produced a near significant overall effect of 4V OT to increase T_IBAT_ when the data were averaged over the 1-h post-injection (T_IBAT 1-hour AVE_) [(F(2,12) = 3.513, *P*=0.063)]. However, there was a significant overall effect of 4V dose to increase T_IBAT_ when the data were averaged over the 2-h post-injection period (T_IBAT 2-hour AVE_) [(F(2,12) = 8.815, *P*=0.006)] (data not shown). Specifically, the high dose of OT (5 μg/μL) was able to stimulate T_IBAT 1-hour AVE_ and T_IBAT 2-hour AVE_ (*P*<0.05) while the lower dose had no significant effect (*P*=NS).

#### Energy intake

Animals used in the telemetry studies were used in these studies. The goal was to confirm that 4V OT reduced energy intake at doses that impacted BAT thermogenesis and activity in DIO rats. Using repeated measures ANOVA, we also found a near significant effect of 4V OT dose to reduce energy intake [(F(8,72) = 2.056, *P* = 0.052] across five time points over the 4-h measurement period. Specifically, there was an overall effect of OT dose to reduce energy intake at 0.5-h post-injection [(F(2,12) = 7.867, *P*=0.007)]. 4V administration of OT at the low dose (1 μg) produced a near significant suppression of 0.5-h energy intake (*P*=0.056) while the high dose (5 μg) reduced 0.5-h energy intake by 78.3±5.4% (*P*<0.05) (**Figure 8D**). 4V administration of OT at the high dose also produced a near significant suppression of energy intake at 1-h post-injection (*P*=0.053).

Overall, these findings indicate that 4V OT increases gross motor activity at doses that stimulate T_IBAT_ and core temperature and reduce energy intake. Furthermore, 4V OT appeared to increase T_IBAT_ prior to core temperature suggesting that OT-elicited changes in T_IBAT_ may also contribute to OT-driven changes in core temperature.

### Study 7B: Determine the extent to which 4V OT activates an early marker of neuronal activation, pERK1/2, within the hindbrain NTS in DIO rats

The goal of this study was to determine if 4V OT treatment is associated with activation of pERK1/2 within hindbrain NTS neurons. Consistent with previous work reporting elevations of pERK1/2-immunoreactive cells at Levels 65–70 in the *Brain Maps 4.0* rat brain atlas [56] in the rat NTS in response to glycemic challenge [64; 75], we identified such elevations in the NTS that mapped to Levels 69 and 70 at the level of the AP (Level 70 data shown in **Figure 9A**). Fifteen minutes after receiving OT into the 4th ventricle, the NTS and DMX display elevated numbers of cells immunoreactive for pERK1/2 (**Figure 9C**). Specifically, phospho-ERK1/2-immunoreactivity (-ir) was observed in fusiform-shaped cells in the NTS, where it also marked slender neurites including what appeared to be axons (*arrowheads* in **Figure 9C**). While further confirmation is needed to determine if these axonal fibers include vagal afferents, pERK1/2 localization to NTS axons is consistent with previous work that identified NTS vagal afferents displaying pERK1/2-ir in response to CCK treatment [63]. The perikaryal NTS labeling we observed appeared to fall within the boundaries of what Swanson has defined as the *caudal subzone (>1840)* of the medial NTS (NTSmc; see Methods for explanation of Swanson’s nomenclature of these regions), although Nissl material from an adjacent tissue series was not always available to help distinguish the boundary zone between the *commissural part (>1840)* (NTSco) and NTSmc at this level. In the DMX, immunoreactivity was observed mainly along the ventral margin of the nucleus in larger, goblet-shaped cells and in smaller, faintly staining triangular-shaped cells (**Figure 9C**). In contrast, control subjects receiving only vehicle solution did not display pERK1/2 immunoreactivity in either brain region (**Figure 9B**). Thus, as with other reported stimuli, 4V OT is also associated with rapid activation of NTS neurons.

**Figure 9A–C:**
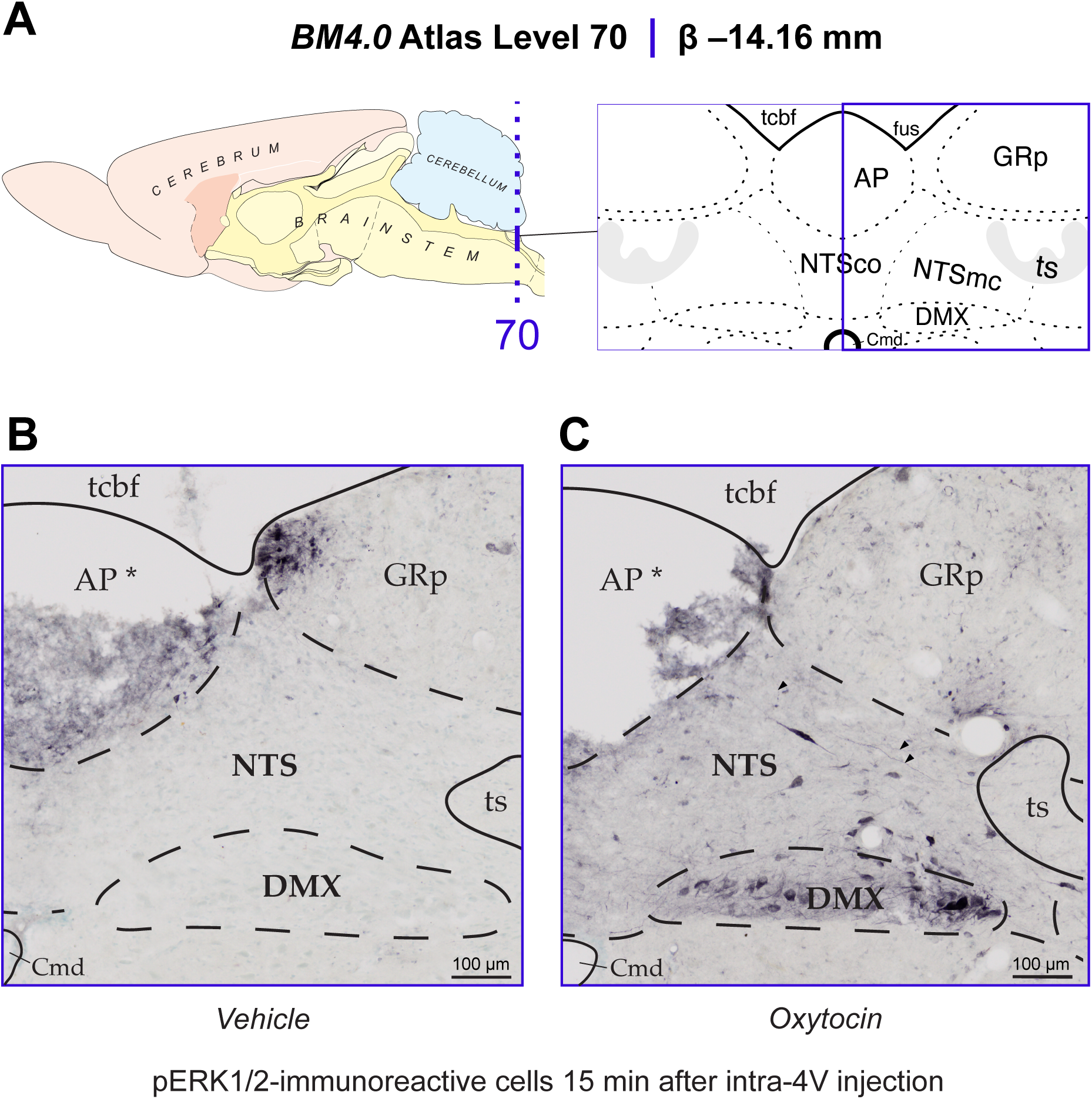
Elevated recruitment of hindbrain nucleus of the solitary tract (NTS) neurons in male DIO rats 15 min after 4V OT administration. *A*, Sagittal view of the rat brain adapted from *BM4.0* depicting the dorsal medullary atlas level area surveyed here. Midline shown in blue. *B/C,* Immunostained hindbrain sections of acute 4V VEH-*(B)* or *(C)* OT-treated subjects. The immunoreactive patterns were used to define the regional boundaries and are represented with dotted lines. Scale bars = 100 µm. Images shown were captured at ✕20 magnification. The header notes the inferred anteroposterior stereotaxic coordinate, expressed as millimeters from the bregma suture on the skull (β). *Note that the area postrema is damaged from histological processing. Abbreviations: AP, *Area postrema (>1840)*; Cmd, *Central canal of medulla (>1840)*; DMX, *Dorsal motor nucleus of vagus nerve (>1840)*; fus, *Funiculus separans (>1840)*; GRp, *gracile nucleus, principal part (>1840)*; NTSco, *Nucleus of solitary tract, commissural part (>1840)*; NTSmc, *Nucleus of solitary tract, medial central part (>1840)*; tcbf, *Transverse cerebellar fissure* [116]; ts, *Solitary tract (>1840)*. All *BM4.0* atlas maps were modified and/or reproduced with permission under the conditions outlined by a Creative Commons BY NC 4.0 license (https://creativecommons.org/licenses/by-nc/4.0/).

## Discussion

The goal of the current studies was to determine if sympathetic innervation of IBAT is required for OT to increase BAT thermogenesis and reduce BW and adiposity in male DIO rats. To assess if OT-elicited increases in BAT thermogenesis require intact SNS outflow to IBAT, we examined the effects of acute 4V OT (1, 5 μg) on T_IBAT_ in DIO rats following sham or bilateral surgical SNS denervation to IBAT. We found that the high dose of 4V OT (5 µg ≈ 4.96 nmol) stimulated T_IBAT_ similarly between sham rats and denervated rats and that the effects of 4V OT to stimulate T_IBAT_ did not require β3-AR signaling. We subsequently determined if OT-elicited reductions of BW and adiposity require intact SNS outflow to IBAT. To accomplish this, we determined the effect of bilateral surgical or sham denervation of IBAT on the ability of chronic 4V OT (16 nmol/day) or vehicle administration to reduce BW, adiposity, and energy intake in DIO rats. We found that chronic 4V OT produced comparable reductions of weight gain, adiposity, leptin levels and energy intake in sham and denervated rats. Lastly, we found that increased activity is not likely to have contributed to the effect of 4V OT to increase both T_IBAT_ and core temperature. 4V OT appeared to increase T_IBAT_ prior to core temperature suggesting that OT-induced elevations in T_IBAT_ may also contribute to OT-induced elevations in core temperature. Collectively, our findings support the hypothesis that sympathetic innervation of IBAT is not a predominant mediator of 4V OT-elicited BAT thermogenesis and reductions of BW and adiposity in male DIO rats.

Our finding showing that the effect of chronic 4V OT to elicit weight loss does not require SNS innervation of IBAT is similar to what we have demonstrated in the mouse model [29]. These findings suggest that OT stimulates IBAT and elicits weight loss through a mechanism that does not require SNS activation of IBAT across rodent species. One outstanding question is how 4V OT activates IBAT if not through SNS outflow to IBAT and signaling through the β3-AR. We found that systemic OT, at a dose that was effective when given into the 4V was ineffective at recapitulating the effects of 4V OT on weight loss and T_IBAT_ in DIO mice [29], suggesting that these effects are not likely a result of 4V OT acting at OTRs in the periphery. One possibility is that 4V OT-elicited stimulation of hindbrain and/or spinal cord OTRs results in the release of epinephrine from the adrenal gland (adrenal medulla) and subsequent stimulation of IBAT. It is possible that epinephrine could be acting through the β1-AR or β2-AR, as both receptor subtypes are expressed in IBAT in both mice [76] and rats [77; 78] (for review see [79]). Others have shown that the β1-AR is important in the control of thermogenesis in rats [80] and mice [81] with the β2-AR receptor potentially playing a more minor role [80] in rodents compared to humans [82; 83; 84], respectively. In addition, L-epinephrine has equal affinity for both β1-AR and β2-AR in CHO cells [85] and it has been shown that epinephrine application to brown adipocytes derived from rat IBAT stimulates respiration and release of fatty acids [86]. While mice that are deficient in epinephrine are still capable of maintaining body temperature in response to cold stress, they are unable to show an elevation of PGC1-alpha or UCP-1 within IBAT, suggesting that epinephrine may be important in mitochondrial uncoupling in IBAT [87]. However, while the hindbrain and spinal cord are a component of a multi-synaptic projection to the adrenal gland [88; 89], only 1% of PVN OT neurons within either the parvocellular or magnocellular PVN were found to have multi-synaptic projections to the adrenal gland [88]. Future studies that examine the effects of 4V OT on 1) plasma levels of epinephrine in sham-operated and IBAT denervated rats and 2) plasma levels of epinephrine, BAT thermogenesis and weight loss in adrenal demedullated rats will be helpful in being able to address the extent to which OT-elicited stimulation of epinephrine from adrenal gland is required for these effects.

We found that IBAT denervation resulted in a significant or near significant reduction of the thermogenic genes, *Dio2*, *UCP-1* and *Prdm16*, from IBAT in DIO rats. These findings are consistent with what others have reported with IBAT UCP-1 protein expression from hamsters [74] and mice [73] following chemical (6-OHDA)-induced denervation of IBAT relative to control animals. Similarly, we and others found a reduction of UCP-1 mRNA expression in IBAT following unilateral or bilateral surgical denervation in mice [29; 70; 71] and hamsters [90], respectively. Our finding showing a reduction of IBAT *Dio2* following IBAT denervation in rats is also consistent with previous findings in mice from our lab and with those of Fischer and colleagues [29; 70].

We also found that chronic 4V OT treatment produced a significant increase of mRNA expression of the thermogenic genes, *Cox8b* and *Adrb1*, in IBAT of sham rats. In addition, 4V OT also produced a near significant increase of mRNA expression of other thermogenic markers (*Gpr120*, *Cidea* and *CN1*) in IBAT in sham rats, some of which were blocked in response to denervation of IBAT (*Cox8b*, *Adrb1*). In contrast, 4V OT increased expression of both thermogenic markers, *Dio2* and *Bmp8b*, in denervated rats but not in sham rats. Similar to the metabolic phenotype of OT [37; 91], OTR [33; 38; 91] deficient mice, both Bmp8b [92] and Dio2 [93; 94] deficient mice have impaired thermogenesis and increased weight gain. However, the obesity phenotype is only observed at thermoneutrality (30°C) but not at 22°C [93] in the Dio2 deficient mice, whereas it occurs at lower temperatures (∼25°C) in the OT, OTR and Bmp8b deficient mice. Whether 4V OT-elicited elevations of Dio2 and Bmp8b mRNA expression in IBAT contributes to the effects of 4V OT to stimulate IBAT thermogenesis in the IBAT denervated rat model will be an important question to address in future studies. While the rats were fasted for 4-h prior to tissue collection to minimize the potential confound of diet-induced thermogenesis on gene expression, it will also be important to include a pair-fed or weight restricted control group to more directly assess the role of energy intake in contributing to these effects. Collectively, these findings raise the possibility that activation of these thermogenic markers might contribute to the effects of 4V OT on BAT thermogenesis in denervated animals.

In addition to changes in IBAT gene expression in response to IBAT denervation, we also found that IBAT denervation was associated with browning of WAT. IBAT denervation increased mRNA expression of the thermogenic genes, *Adrb3* and *UCP-1*, in EWAT (indicative of browning of WAT). However, our finding showing an elevation of EWAT UCP-1 in response to IBAT denervation in rats is in contrast to the findings from Nguyen, who failed to find any change in UCP-1 protein in EWAT between sham-operated and IBAT-denervated hamsters [74]. These different outcomes may potentially be explained by species differences. While UCP-1 mRNA expression from IWAT was not examined in our study, the findings pertaining to EWAT UCP-1 gene expression in surgically denervated rats are consistent with findings from other white adipose tissues, namely IWAT, which show an elevation of IWAT UCP-1 protein in both hamsters [74] and mice [73] following chemical (6-OHDA)-induced denervation of IBAT relative to control animals. Another study found what appeared to be a marked increase in browning of IWAT (as indicated by UCP-1 immunohistochemistry) following bilateral surgical denervation of IBAT in mice but the data were not quantified [72]. In addition, Nguyen also found that IBAT denervation resulted in an elevation of IWAT temperature (functional readout of WAT thermogenesis) in response to cold stress relative to sham animals [74]. 4V OT resulted in increased expression of *Adrb3* and *Gpr120* as well as decreased expression of CN1 in EWAT of sham-operated rats while it also increased UCP-3 mRNA expression in denervated rats. It remains to be determined whether different thermogenic markers in EWAT contribute, in part, to the metabolic effects of 4V OT in sham (*Adrb3*, *Gpr120*, *CN1*) and denervated rats (*UCP-3*).

One of the remaining questions from this study is whether the changes in thermogenic gene expression in other WAT depots in response to IBAT denervation may impact the effectiveness of OT to stimulate BAT thermogenesis and elicit weight loss. Previous findings have identified both distinct and overlapping CNS circuits that regulate SNS outflow to IBAT and IWAT [74]. Namely, OT neurons within the parvocellular PVN send multi-synaptic projections to IBAT [95; 96], EWAT [97; 98] and IWAT [95; 97]. A small subset of OT neurons within the parvocellular PVN also overlap and project to both IBAT and IWAT [95]. Thus, PVN OT neurons are anatomically positioned to impact SNS outflow to IBAT and IWAT and increase both BAT and WAT thermogenesis, respectively. It is not clear if these effects are mediated by the same OT neurons or through distinct OT neurons that project to the hindbrain NTS [99; 100] and/or spinal cord [100], both of which are areas that can alter SNS outflow to IBAT and BAT thermogenesis [101; 102]. Recent findings from Ong reported that viral-elicited knockdown of OTR mRNA within the hindbrain dorsal vagal complex failed to block the effects of 4V OT to stimulate core temperature [103], suggesting that OTRs within other hindbrain sites and/or spinal cord may be important in mediating these effects in rats. While our pERK1/2 analysis was confined to the dorsal vagal complex, it is possible that 4V OT could have reached OTR in other brain sites that can impact BAT thermogenesis, as reported in studies to occur in the spinal cord [30; 104] and/or rostral raphe pallidus [33] (which corresponds to *pallidal raphe nucleus >1840*) (RPA) according to Swanson’s nomenclature [61]. Thus, future studies should examine the extent to which OTRs in regions of the hindbrain NTS not targeted by Ong and colleagues [103], as well as other hindbrain sites such as the RPA [33; 102; 105; 106; 107] or spinal cord [30], contribute to the effect of 4V OT on BAT thermogenesis.

Our findings suggest that increased locomotor activity is not likely to be the main driver of 4V OT on T_IBAT_. 4V OT-elicited changes in both T_IBAT_ and core temperature preceded significant changes in gross motor activity, as did the phospho-ERK1/2 activation patterns, which we observed in the NTS within 15 minutes of 4V OT administration. It is possible that changes in gross motor activity in response to the higher dose of OT (5 μg) could be potentially important in contributing or maintaining the increase in T_IBAT_ and core temperature at 1.5-h post-injection, but this does not appear to be the case at the earlier time points. Consistent with our findings linking CNS OT to activity, Sakamoto and colleagues reported that intracerebroventricular (ICV) administration of a lower dose of OT (0.5 μg) increased activity in mice [108]. In addition, Noble and colleagues found that localized administration of OT (1 nmol ≈ 1.0072 μg) into the ventromedial hypothalamus also increased physical activity at 1-h post-injection in rats [13]. Sutton and colleagues also reported that chemogenetic activation of PVN OT neurons produced an elevation in locomotor activity, energy expenditure and subcutaneous IBAT temperature [30] in *Oxytocin-Ires Cre* mice. This finding raises the possibility that release of endogenous OT following activation of PVN OT neurons may also elicit increases in activity. While our limited sample size and restraint stress (see below) might have hampered our ability to measure more robust effects of 4V OT on activity at earlier time points, our findings suggest that locomotor activity does not contribute to the effects of 4V OT on T_IBAT_ in DIO rats.

Similar to what we observed with our recent studies in mice, one limitation of our behavioral studies is that handling or restraint stress may have limited our ability to observe more pronounced differences between drug and vehicle on T_IBAT_ [109] and activity during the time period when the effect of the drug on T_IBAT_ or activity is relatively short-lived or small. As a way to minimize the effect of restraint stress on T_IBAT_, we adapted the rats to handling and mock injections one week prior to the experiment and by administering the drugs during the early part of the light cycle when T_IBAT_ [53] and catecholamine levels are lower [54]. While the effects of CL 316243 continued well beyond the short-lived effects of restraint/vehicle stress on T_IBAT_ in our rat studies (∼30-45 minutes), the effects of OT and SR 59230A in **Study 5** were relatively short-lived and may have been masked by restraint stress. Thus, stress-induced epinephrine, from the adrenal medulla [110; 111; 112], may have activated β3-AR in IBAT to stimulate T_IBAT_, even in IBAT-denervated rats.

We acknowledge that other BAT depots (axillary, cervical, mediastinal and perirenal depots), all of which show cold-induced elevations of UCP-1 [113], might act to have contributed, in part, to the effects of 4V OT to elicit weight loss in IBAT-denervated rats. We maintained the focus on IBAT given that this depot is the best characterized of all BAT depots [114]. However, IBAT is thought to contribute to ≥ 70% of total BAT mass [115] or approximately 45% of total thermogenic capacity of BAT [70]. Moreover, we acknowledge the potential contribution of WAT thermogenesis to OT-elicited weight loss and BAT thermogenesis, particularly EWAT, given that chronic 4V OT was able to elevate EWAT UCP-3 in IBAT of denervated rats. As indicated above, there is crosstalk between SNS circuits that innervate WAT and IBAT [74]. In addition, there is increased NE turnover and IWAT UCP-1 mRNA expression in IBAT-denervated hamsters [74]. It will be important to 1) confirm our EWAT gene expression findings and to include an analysis of IWAT in IBAT-denervated rats and 2) develop a model to better understand whether the other BAT and specific WAT depots contribute to 4V OT-elicited weight loss.

In summary, our findings demonstrate that acute 4V OT (5 µg) produced comparable increases in T_IBAT_ in both denervated and sham rats. We subsequently found that chronic 4V OT produced similar reductions of BW gain and adiposity in both sham and denervated rats. Importantly, our findings suggest that there is no change or obvious functional impairment in the response of the β3-AR agonist, CL 316243, to activate IBAT in rats with impaired SNS innervation of IBAT in comparison to sham-operated rats. Furthermore, we acknowledge that increased activity is not likely to have contributed to the effect of 4V OT to increase both T_IBAT_ and core temperature. In addition, 4V OT appeared to increase T_IBAT_ prior to core temperature suggesting that OT-induced elevations in T_IBAT_ may also contribute to OT-induced elevations in core temperature. Together, these findings support the hypothesis that sympathetic innervation of IBAT is not a predominant mediator of OT-elicited increases in BAT thermogenesis and reductions of BW and adiposity in male DIO rats.

## ACKNOWLEDGMENTS

The authors thank the technical support of Nishi Ivanov, Hailey Chadwick, Alex Vu and Ana R. Arvizu Morales. In addition, the authors are appreciative of the efforts by Drs. Michael Schwartz and Dianne Lattemann for providing feedback throughout the course of these studies. Disclosures: JEB had a financial interest in OXT Therapeutics, Inc., a company developing highly specific and stable analogs of oxytocin to treat obesity and metabolic disease. The author’s interests were reviewed and are managed by their local institutions in accordance with their conflict-of-interest policies. The other authors have nothing to report.

**Supplemental Study 1: Determine if surgical denervation of IBAT blocked the ability of 4V OT to increase T_IBAT_ in lean rats.** Rats (N= 6 at study onset) were used for this study. The goal of this study was to determine if OT-elicited increase in T_IBAT_ requires intact SNS outflow to IBAT in lean rats. By design, rats were lean as determined by both BW (511.6 ± 2.4 g) and adiposity (3.3 ± 0.3 g fat mass; 18.1 ± 11.1% adiposity) after maintenance on the chow (13% kcal from fat; N=6/group) for approximately 1 week prior to denervation procedures and implantation of temperature transponders underneath IBAT. Rats were otherwise treated identically to those used in **Study 4**.

**Supplemental Study 1:**

Tissue samples from **Supplemental Study 1** were screened for NE content and compared with a previous cohort of sham-operated rats from **Study 1**. However, there was no age-matched cohort to use for comparison purposes to calculate reduction of IBAT NE content as the cohort used in **Supplemental Study 1** was heavier than the sham-operated cohort used in **Study 1** (354 ± 4.6 g) [(F(1,9) = 608.553, *P*<0.001]. IBAT NE content from sham-operated animals was only used to screen whether IBAT NE content was above the acceptable threshold for a successful IBAT denervation. One of the six animals was removed on account of having a failed IBAT denervation procedure NE content was reduced by 93.8 ±2.6% in IBAT from denervated rats relative to IBAT from sham-operated control rats [(F(1,9) = 16.390, *P*<0.01]. IWAT NE content was also reduced in the denervated group [(F(1,9) = 7.411, *P*<0.05)] but this might have been due, in part, to the differences in age of the animals between cohorts. In contrast, NE content was unchanged in pancreas but elevated in liver [(F(1,9) = 8.684, *P*<0.05] in denervated rats relative to sham rats.

In denervated rats, 4V OT (5 μg/μL) increased T_IBAT_ at 0.5, 0.75, 1, 1.25, 1.5 and 1.75-h post-injection (*P*<0.05; **Supplemental Figure 1A**). The high dose (5 μg/μL) also stimulated T_IBAT_ when measuring change in T_IBAT_ relative to baseline T_IBAT_ at 0.5, 0.75, 1, 1.25, 1.5, 1.75 and 2-h post-injection (*P*<0.05; **Supplemental Figure 1B**). The low dose (1 μg/μL) increased T_IBAT_ at 1.5-h post-injection (*P*<0.05; **Supplemental Figure 1A**). The low dose (1 μg/μL) produced a near significant stimulation of T_IBAT_ when measuring change in T_IBAT_ relative to baseline T_IBAT_ at 1- and 1.5-h post-injection (0.05<*P*<0.1; Supplemental Figure 1B).

**Supplemental Study 2: Determine the extent to which β3-AR-induced activation of IBAT requires activation of β3-AR to increase T_IBAT_ in DIO rats.**

Rats (N= 19 at study onset) were used for this study. Rats were fed *ad libitum* and maintained on HFD for approximately 4.5 months prior to receiving 4V cannulas and/or temperature transponders [28]. Rats were allowed to recover for at least 4 weeks during which time they were adapted to a daily 4-h fast, handling, and mock injections.

The goal of this study was to determine a dose of the β3-AR antagonist that was sufficient to block the effects of the β3-AR agonist on T_IBAT_ in DIO rats. The dose of the β3-AR antagonist would then be used in **Study 6** to determine if 4V OT required β3-AR signaling to stimulate T_IBAT_ in DIO rats. By design, DIO rats were obese as determined by both BW (712 ± 23.1 g) and adiposity (217 ± 12.1 g fat mass; 31 ± 1.0% adiposity) after maintenance on the HFD for 4.5 months prior to sham/denervation procedures.

In the absence of the β3-AR antagonist, the β3-AR agonist increased T_IBAT_ at 0.25, 0.5, 0.75, 1, 1.5, 0.75, 2 and 4-h post-injection (*P*<0.05) and produced a near significant stimulation of T_IBAT_ at 180-min post-injection (0.05<*P*<0.1) (**Supplemental Figure 2**). These effects were blocked at 0.5-h post-injection of the β3-AR agonist. Together, these findings confirm the specificity of a β3-AR mediated effect of the β3-AR agonist on T_IBAT_ and identified a dose of the β3-AR antagonist that was sufficient to block the effects of the β3-AR agonist to stimulate T_IBAT_.

Two-way repeated-measures ANOVA revealed a significant main effect of β3-AR agonist to increase T_IBAT_ at 0.5-h post-injection [F(1,48) = 5.844, *P*=0.019], a significant main effect of the β3-AR antagonist on T_IBAT_ at 0.5-h post-injection [F(1,48) = 5.671, *P*=0.021] and a significant interaction between the β3-AR agonist and the β3-AR antagonist on T_IBAT_ at 0.5-h post-injection [F(1,48) = 5.728, *P*=0.021].

Overall, these findings confirmed the specificity of a β3-AR mediated effect of the β3-AR agonist on T_IBAT_ and identified a dose of the β3-AR antagonist that was sufficient to block the effects of the β3-AR agonist to stimulate T_IBAT_.

**Supplemental Study 3: Determine the extent to which OT-induced activation of sympathetic outflow to IBAT contributes to its ability to elicit weight loss in rats with more pronounced diet-induced obesity.**

Rats (N= 20 at study onset) were used for this study. Rats were fed *ad libitum* and maintained on HFD for approximately 8.75 months prior to receiving 4V cannulas and minipumps to infuse vehicle or OT (16 nmol/day) over 29 days as previously described [28]. Daily energy intake and BW were also tracked for 29 days. Animals were euthanized by rapid conscious decapitation at 9 weeks post-sham or denervation procedure. Trunk blood and tissues (IBAT, EWAT, IWAT, liver and pancreas) were collected from 4-h fasted rats and tissues were subsequently analyzed for IBAT NE content to confirm success of denervation procedure relative to sham operated animals and other tissues (EWAT, IWAT, liver and pancreas).

The goal of this study was to determine if OT-elicited weight loss requires intact SNS outflow to IBAT. By design, DIO rats were obese as determined by both BW (790.7 ± 23.8 g) and adiposity (281.9 ± 16.3 g fat mass; 35.3 ± 0.1.3% adiposity) after maintenance on the HFD for 8.75 months prior to sham/denervation procedures.

All IBAT tissues from **Supplemental Study 3** animals were analyzed for IBAT NE content and one of the ten animals were removed on account of having a failed IBAT denervation procedure IBAT NE content was reduced in denervated rats by 84.8±4.3% in denervated rats relative to sham-operated control rats [(F(1,16) = 128.544, *P*=0.000)]. In contrast, NE content was unchanged in IWAT, EWAT or pancreas in denervated rats relative to sham rats (*P*=NS). In contrast, there was a reduction of NE content in liver in denervated rats relative to sham rats [(F(1,16) = 4.654, *P*=0.047)]. There was no significant difference in BW between sham and denervation groups at the end of the study (*P*=NS; data not shown).

In sham-operated rats, as expected, 4V vehicle resulted in 4.3 ± 1.0% weight gain relative to vehicle pre-treatment [(F(1,4) = 15.268, *P*=0.017]. In contrast, 4V OT reduced BW by 4.2 ± 1.2% relative to OT pre-treatment [(F(1,4) = 10.931, *P*=0.030] (**Supplemental Figure 3A**) and it also reduced weight gain throughout the 29-day infusion period (**Supplemental Figure 3B**). OT treatment reduced weight gain on days 4-29 (*P*<0.05) and produced a near significant reduction of weight gain on day 3 (*P*=0.068). 4V OT reduced relative fat mass (pre- vs post-intervention; **Supplemental Figure 3C**) (*P*<0.05) and produced a reduction in the relative lean body mass (pre- vs post-intervention; *P*<0.05) without impacting total fat mass or lean body mass (*P*=NS). These effects that were mediated, at least in part, by a modest reduction of energy intake that persisted throughout the first two weeks of the treatment period (**Supplemental Figure 3D**; *P*<0.05). 4V OT also produced a near significant reduction of energy intake during weeks 3 and 4 of the treatment period (**Supplemental Figure 3D**; 0.05<*P*<0.1).

In denervated rats, as expected, 4V vehicle resulted in 4.7 ± 0.8% weight gain relative to vehicle pre-treatment [(F(1,3) = 44.734, *P*=0.007] (**Supplemental Figure 3A**). In contrast, 4V OT failed to reduce BW relative to OT pre-treatment but it reduced weight gain throughout the 29-day infusion period (**Supplemental Figure 3B**). OT treatment reduced weight gain on days 4-29 (*P*<0.05) and it produced a near significant reduction of weight gain on day 3 (*P*=0.05). 4V OT reduced relative fat mass (pre- vs post-intervention; **Supplemental Figure 3C**) (*P*<0.05) without effecting total fat mass or lean body mass (*P*=NS). These effects that were mediated, at least in part, by a modest reduction of energy intake that was evident during weeks 1 and 2 of the treatment period (**Supplemental Figure 3D**; *P*<0.05).

There was no significant difference in BW between sham and denervation groups at the end of the study (≈ 9-weeks post-sham/denervation surgery) (*P*=NS; data not shown).

Two-way ANOVA revealed a significant main effect of OT to reduce body weight gain on day 29 [F(1,14) = 39.841, *P*<0.01] but no overall effect of denervation [F(1,14) = 0.292, *P*=0.598] or an interactive effect between OT and denervation on body weight gain on day 29 [F(1,14) = 0.094, *P*=0.908]. Two-way ANOVA revealed no significant main effect of OT [F(1,14) = 1.278, *P*=0.277] or denervation on fat mass [F(1,14) = 0.051, *P*=0.825] or an interactive effect between OT and denervation on fat mass [F(1,14) = 0.443, *P*=0.517].

Overall, these findings demonstrate that sympathetic innervation of IBAT is not a predominant mediator of 4V OT-elicited reductions of BW and adiposity in male DIO rats.

## Figure legend

**Supplemental Figure 1A–D:**
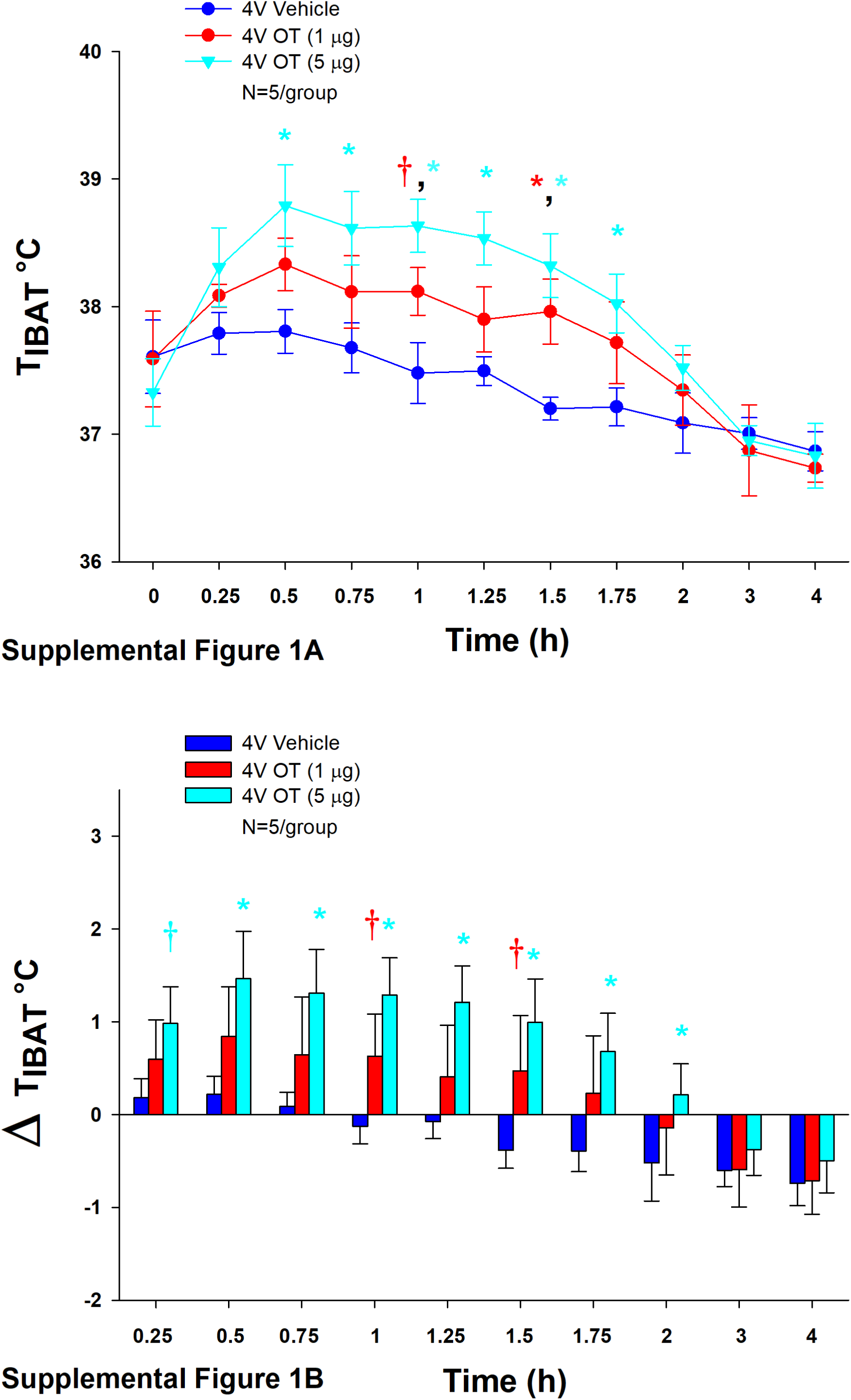
Effect of 4V OT on IBAT temperature (T_IBAT_) in lean rats. Rats (N=6 total) were fed *ad libitum* and maintained on low fat chow diet (13% kcal from fat) (N=6/group) for approximately 1 week prior to underdoing SNS denervation procedures and implantation of temperature transponders underneath the left IBAT depot. Rats subsequently received 4V cannulations. Rats were allowed to recover for at least 2 weeks during which time they were adapted to a daily 4-h fast, handling, and mock injections. Rats subsequently received injections of 4V OT (1 or 5 μg/μL) or vehicle where each animal received each treatment at least 48-h intervals. *A*, Effect of 4V OT on T_IBAT_ in IBAT denervated lean rats; *B*, Effect of 4V OT on change in T_IBAT_ relative to baseline T_IBAT_ (delta T_IBAT_) in IBAT denervated lean rats. Data are expressed as mean ± SEM. \**P*<0.05, †0.05<*P*<0.1 4V OT vs. vehicle.

**Supplemental Figure 2A–B:**
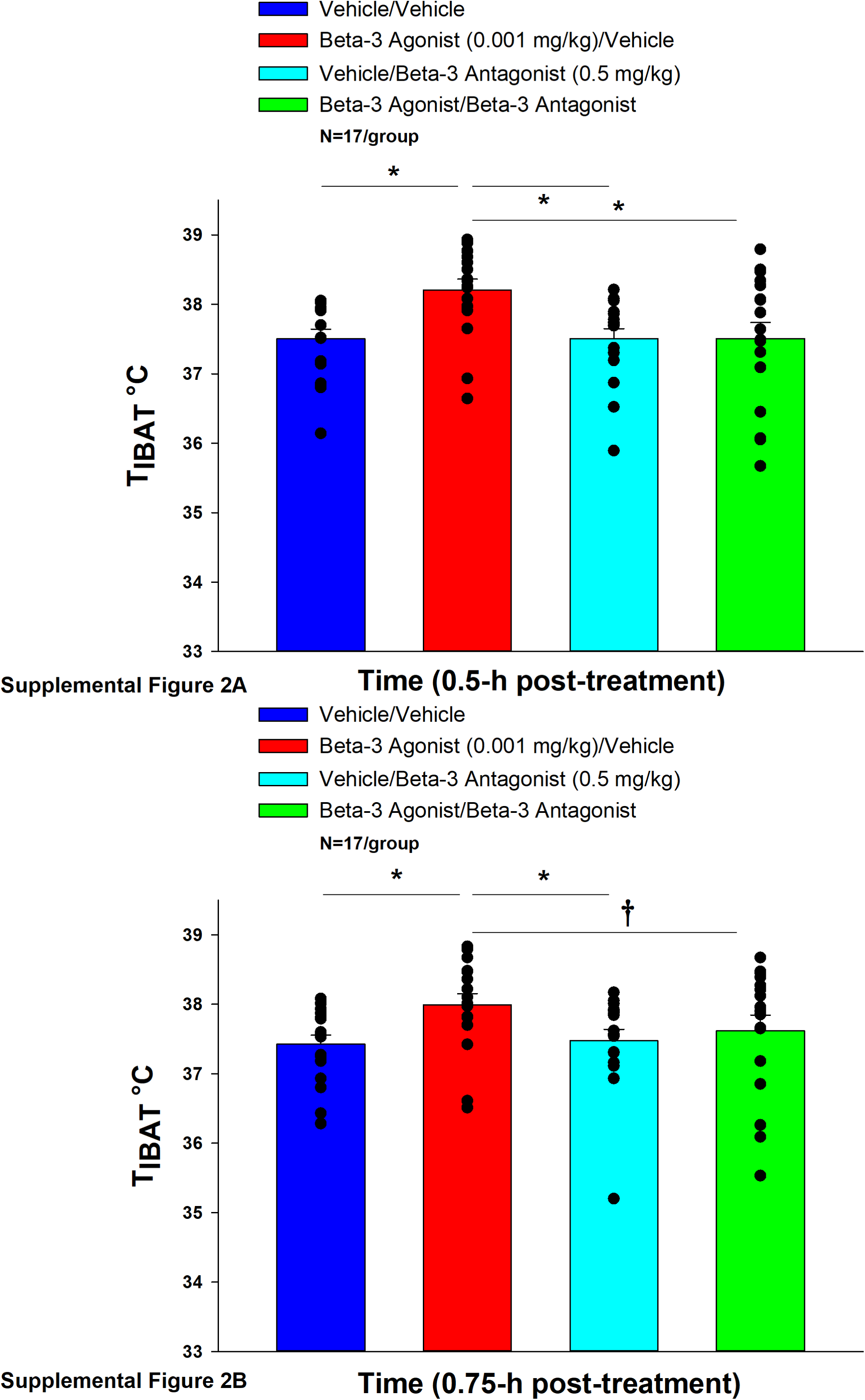
Effect of β3-AR antagonist, SR59230A, on the ability of the β3-AR agonist, CL 316243, to increase T_IBAT_ in DIO rats. Rats (N=17 total) were maintained on HFD (60% kcal from fat; N=17/group) for approximately 4.25 months prior to being implanted with temperature transponders underneath IBAT. Animals were subsequently adapted to a 4-h fast prior to receiving acute IP injections of the β3-AR agonist, SR59230A or vehicle approximately 20 minutes prior to IP injections of the β3-AR agonist, CL 316243. *A*, Effect of the β3-AR antagonist (SR-59230A) pre-treatment on the ability of the β3-AR agonist, CL 316243, to stimulate T_IBAT_ at 0.5-h post-injection; *B*, Effect of the β3-AR antagonist (SR-59230A) pre-treatment on the ability of the β3-AR agonist, CL 316243, to stimulate T_IBAT_ at 0.75-h post-injection. Data are expressed as mean ± SEM. \**P*<0.05, †0.05<*P*<0.1 OT vs. vehicle.

**Supplemental Figure 3A–D:**
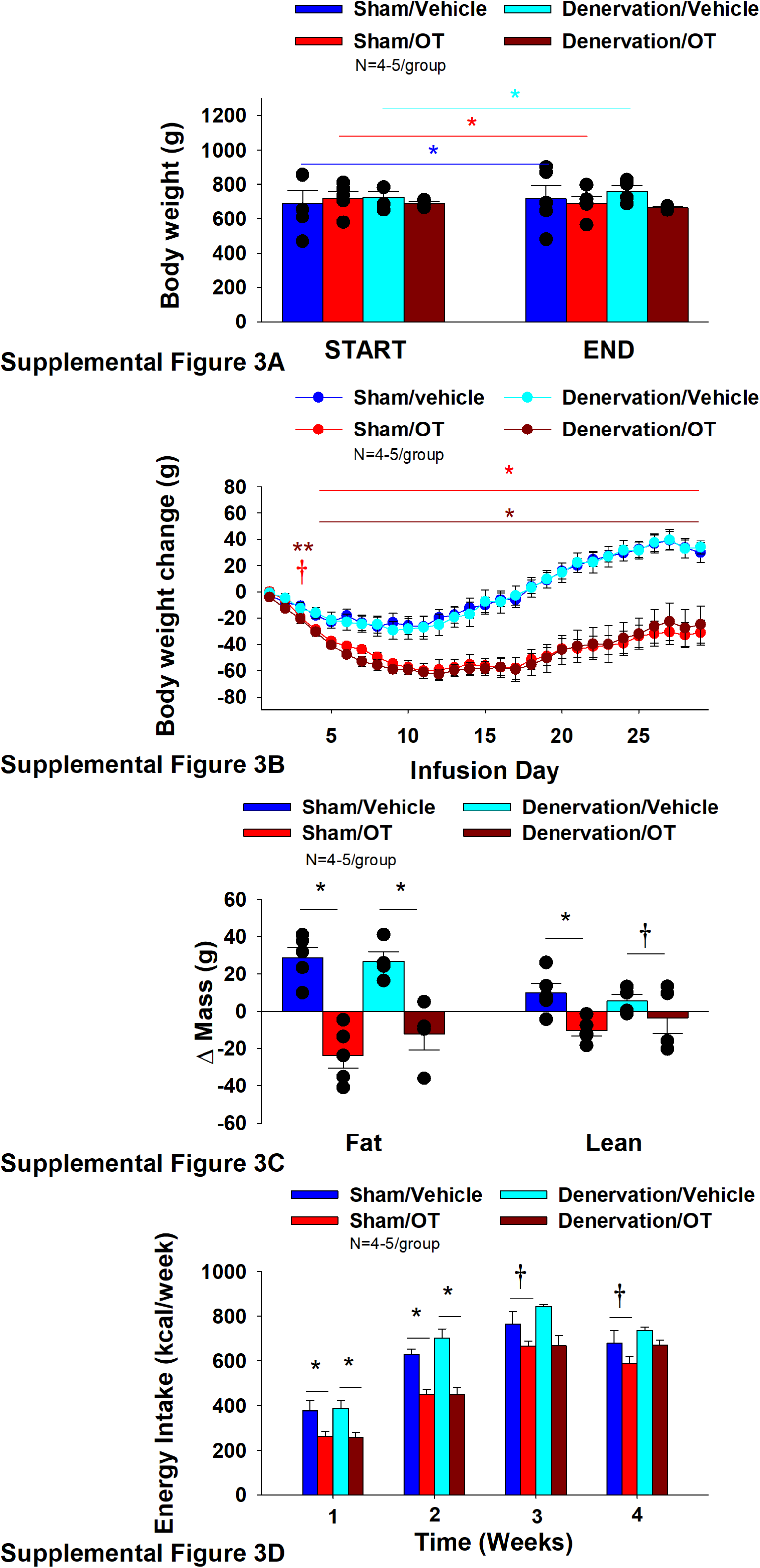
Effect of chronic 4V OT infusions (16 nmol/day) on BW, adiposity, and energy intake post-sham or IBAT denervation in male DIO rats with more pronounced diet-induced obesity. *A*, Rats (N=18 total) were maintained on HFD (60% kcal from fat; N=4-5/group) for approximately 8.75 months prior to undergoing a sham or bilateral surgical IBAT denervation. Rats were subsequently implanted with 4V cannulas and allowed to recover for 2 weeks prior to being implanted with subcutaneous minipumps that were subsequently attached to the 4V cannula. *A*, Effect of chronic 4V OT or vehicle on BW in sham operated or IBAT denervated DIO rats; *B*, Effect of chronic 4V OT or vehicle on BW change in sham operated or IBAT denervated DIO rats; *C*, Effect of chronic 4V OT or vehicle on adiposity in sham operated or IBAT denervated DIO rats; *D*, Effect of chronic 4V OT or vehicle on adiposity in sham operated or IBAT denervated DIO rats. Data are expressed as mean ± SEM. \**P*<0.05, †0.05<*P*<0.1 OT vs. vehicle.

